# Volatile social environments can favour investments in quality over quantity of social relationships

**DOI:** 10.1101/2021.10.01.462760

**Authors:** Thomas G. Aubier, Hanna Kokko

## Abstract

Cooperation does not occur in a vacuum: interactions develop over time in social groups that undergo demographic changes. Intuition suggests that stable social environments favour developing few but strong reciprocal relationships (a ‘focused’ strategy), while volatile social environments favour the opposite: more but weaker social relationships (a ‘diversifying’ strategy). We model reciprocal investments under a quality-quantity tradeoff for social relationships. We find that volatility, counterintuitively, can favour a focused strategy. This result becomes explicable through applying the theory of antagonistic pleiotropy, originally developed for senescence, to social life. Diversifying strategies show superior performance later in life, but with costs paid at young ages while the social network is slowly being built. Under volatile environments, many individuals die before reaching sufficiently old ages to reap the benefits. Social strategies that do well early in life are then favoured: a focused strategy leads individuals to form their first few social bonds quickly and to make strong use of existing bonds. Our model highlights the importance of pleiotropy and population age structure for the evolution of cooperative strategies and other social traits, and shows that it is not sufficient to reflect on the fate of survivors only, when evaluating the benefits of social strategies.

## INTRODUCTION

The evolution of cooperation is a central theme in biology (West et al. 2007, 2021). Past theoretical work has elucidated how cooperative traits can be advantageous over non-cooperative traits that better exploit the public good of cooperation. While inclusive fitness theory explains why cooperation among kin can evolve (Hamilton 1964, Hamilton 1970, Bourke 2011, Kay et al. 2020), reciprocity theory explains how cooperation between individuals that are not necessarily related can be enforced by reward and punishment (Trivers 1971, Axelrod & Hamilton 1981, Clutton-Brock 2009). In animal societies, such reciprocal cooperations can lead to the build-up of social partnerships, as shown in humans (Axelrod & Hamilton 1981, Rand et al. 2011), vampire bats (Wilkinson 1984, Carter & Wilkinson 2013, Carter et al. 2017), some nonhuman primates (Packer 1977, Lehmann et al. 2007, Schino and Aureli 2008), some fishes (Fischer 1984, Hart 2016) and some birds (McDonald 2007, Dakin & Ryder 2020).

A gap exists between simple evolutionary models of cooperation and the complex psychology of helping decisions in animal societies (Connor 2010, Cantor et al. 2021). In particular, Connor (2010) argue that thinking ‘beyond the dyad’ (i.e., not considering solely fixed pairs of individuals) is necessary if we are to understand the evolution of cooperative strategies in animal societies (as assumed when taking the ‘biological market’ perspective; Noë & Hammerstein, 1995). For animals that live in complex societies, much cooperation takes place in the context of stable social bonds that seem analogous to human friendships. Yet, very few theoretical studies have investigated the evolution of cooperative strategies within an explicit social network, and have considered how the network itself evolves as a result of individual interactions (Ohtsuki et al. 2006, Delton et al. 2011, Dakin & Ryder 2020, Akçay 2020).

A particularly intriguing question is what cooperative strategy evolves when individuals face an unavoidable tradeoff between quantity and quality of their social relationships. Cognitive limitations (Dunbar 1992, Dunbar 1993, Tamarit et al. 2018), and time or other resource budgets (Ozaktas 1996), create constraints such that one cannot possibly be ‘everyone’s best friend’. Differential investment in quantity vs. quality of social relationships has been observed in humans (Sutcliffe & Crabbe 1963, Amato 1993, Entwisle et al. 2007, Sato & Zenou 2015, Vacca 2019), in nonhuman primates (Silk et al. 1999, Ellis et al. 2019, Morrison et al. 2020), and in other mammals such as kangaroos (Menz et al. 2020) and giraffes (Bond et al. 2021). Carter et al. (2017) showed that cooperative vampire bats investing in quantity of social relationships (with unrelated individuals) at the expense of relationship quality cope better with a volatile social environment; the authors named this strategy a ‘social bet-hedging strategy’, as the diversified portfolio of acquaintances protects against the worst-case outcome of none of one’s few friends remaining alive. Nonetheless, there is surprisingly little theory on the evolutionary forces shaping the cooperative strategies that ultimately determine the individuals’ social environment, which has potentially important consequences on survival (Snyder-Mackler 2020), and which in turn may feed back into the stability of the social environment itself.

Here, we model the competition between focused strategies (our shorthand for the tendency to form few but strong relationships) and diversifying strategies (where individuals readily develop social ties with strangers but, due to the quality-quantity trade-off, these remain weaker; Carter et al. 2017, Cantor et al. 2021) along a continuum where the readiness to invest in new social bonds varies. Perhaps counterintuitively, we show that volatility in the social environment can favour a focused, rather than a diversifying, approach. The result becomes explicable when realizing that the ultimate performance of the eventual network is not the sole criterion for its evolutionary success. The diversifying approach builds an excellent network but does so slowly, and the focused strategy outperforms it during early life. If many individuals are relatively short-lived (due to volatility), it is better to sacrifice late-life performance for improved success in early life. This shows that the antagonistic pleiotropy theory of senescence (Williams 1957, recent review: Flatt & Partridge 2018) is of relevance for the development of social strategies, not only for somatic maintenance.

## THE MODEL

### Overview

We model individuals that live in groups of N individuals. Group size is kept constant by recruitment of new group members as soon as an existing member has died.

All individuals share resources and thereby participate in the dynamics of social bond forming over their entire lifetime. Reproduction is asexual for simplicity, and generations are overlapping. While all individuals use the same structural rules of Bayesian updating of social bonds (details below), they differ in their *a priori* propensity to establish contact with individuals with whom they do not have an interaction history yet. For brevity we call such individuals ‘strangers’, though note that they live permanently in the same group. For humans, a classroom provides the appropriate mental image for this type of stranger: while all children share the classmate status, each child will only name a subset of all classmates as friends, and ‘strangers’ in this context are those who, despite physical proximity, are not relevant for a focal child’s social network. Obviously, real classroom settings can be more complex, with agonistic interactions (e.g. bullying) also present; we do not imply that bonds cover all such cases, instead we invite the reader to use this image simply as a reminder that strangers, as defined by us, are not *per se* unavailable due to distance.

These propensities to establish contact with strangers are genetically encoded, with two independent traits that impact an individual *i*‘s propensity to (1) ask resources from strangers (A_i_) or (2) to give resources to strangers (G_i_). High values indicate a tendency to diversify one’s social relationship portfolio, low values indicate a focused approach to relationships. There is both resource-independent and resource-dependent mortality, and each vacancy created by death is replaced by a new recruit, with the parent chosen randomly from the population of living individuals. Selection on traits operates based on survival: individuals whose traits give them a reliable resource supply contribute disproportionately to future generations by virtue of them living longer. Recruits inherit their (single) parent’s trait with some mutation, allowing A_i_ and G_i_ to evolve, with the social network also changing as an emergent property of the population.

Inspired by vampire bat biology (Wilkinson 1984, Carter & Wilkinson 2013, Carter et al. 2017), we assume that each individual attempts to perform independent foraging at each time step, but this may fail, creating a constant supply of successful (satiated) and unsuccessful (needy) individuals who thereafter can interact socially, allowing resources to be donated to unsuccessful individuals. Note that we assume pre-existing willingness to help others (we do not give individuals the option to cheat), as our model is designed to investigate the evolution of focused versus diversified relationship tendencies, and the resultant narrow or broad social networks, rather than the origins of cooperation *per se*. For the same reason, in our main analysis, we also ignore complications brought about by kin recognition and preferential helping among kin; we thus do not track relatedness of the individuals (but see additional simulations in Figs. S1 and S2).

Cooperative traits, A_i_ and G_i_, jointly determine to what extent individuals focus vs. diversify cooperative investments. We focus on the implications of the volatility of the social environment on the evolution of those cooperative traits. We explicitly implement four sources of volatility. First, memory of past interactions between each pair of individuals can become erased with probability P_erase_, making the individuals strangers to each other. Second, individuals may be unsuccessful during foraging with probability P_unsucc_. Third, successful individuals may be unavailable with probability P_unav_, not being able to give any resources. Fourth, individuals may randomly die with probability P_die_. Increasing each of these probabilities associates with an increased volatility of the social environment, with strong social bonds being lost temporarily or permanently.

### Social bonds

#### Basic properties

We model the strength of a social bond between each pair of individuals as a continuous variable (ϵ [0,1]) that changes when individuals interact with each other. We first list some desirable properties of how to model the social bond, before proceeding to the mathematical definitions. The bond should be strong if individuals have helped each others reciprocally in the past (allowing us to call them ‘partners’, without however implying that the relationship excludes having outside options). Strong bonds mean that an individual that has helped another individual is predisposed to ask resources from its partner should the need arise. Conversely, an individual that has been helped by another individual will tend to return the favour. As a flipside, an individual refusing to help another individual is less tempted to ask resources from this same individual, who, as a mirror image of the argument, will be little inclined to help.

#### Perspective-dependence of the strength of social bonds

Each individual uses Bayesian updating to formulate its own estimate of the social bond. Individuals with different trait values, A_i_ and G_i_, have different estimates of the strength of the same social bond. In particular, individuals with high A_i_ and G_i_ trait values have high estimate values of the baseline strength of social bonds with strangers, and are therefore more inclined to interact with them than individuals with low A_i_ and G_i_ trait values do. Hence, we define the strength of social bonds from the perspective of each individual, with **∏**_i_(i,j) representing the estimate by individual *i* of the strength of the social bond between individuals *i* and *j*, and **∏**^j^(i,j) representing the estimate by individual *j* of this same quantity. These estimates directly determine the cooperative behaviour of individuals (see subsection ‘Resource donation’ below). To describe the properties of the emerging social network, however, we act as an unbiased external observer who gives the same weight to the perspective of each actor, as detailed later.

#### Estimates of the strength of social bonds

We now come to the expressions of **∏**^i^(i,j) and **∏**^j^(i,j), i.e., the estimates of the strength of the social bond between individuals *i* and *j* if we take the perspective of individual *i* and *j*, respectively. For individual *i*, **∏**^i^(i,j) corresponds to the product of two quantities, **∏**^i^(i,j) = π^i^_j→i_ x π^i^_i→j_, presented below (superscript *i* refers to the perspective of individual *i*). Likewise, for individual *j*, **∏**^j^(i,j) = π^j^_j→i_ x π^j^_i→j_.

Individual *i* ‘s estimate of individual *j* ‘s trustworthiness is π^i^_j→i_. It is calculated as a Bayesian estimate (belief) of the probability that individual *j* donates resources when individual *i* has placed a request. Hence, superscript *i* of π^i^_j→i_ indicates that this is individual *i*’ Bayesian estimate, and the arrow of subscript ‘j→i’ of π^i^_j→i_ indicates who donates resources to whom when calculating the Bayesian estimate. Consider first the standard case, where some interactions have already taken place; the case with *j* being a stranger is dealt with afterwards. Each past interaction (defined as the outcome when individual *i*has placed a request, with multiple requests in one time step counting as separate interactions) has resulted in individual *j*either giving a resource package or not (see subsection ‘Resource donations’ below). The number of resource donations from all interactions that have occurred between *i* and *j* so far follows a binomial distribution. The conjugate distribution for a binomial is a Beta distribution, so specifying the prior probability distribution (belief before the next interaction) as a Beta distribution allows us to retain the Beta distribution in the posterior (the updated belief after an interaction). Specifically, we assume the prior follows a Beta distribution B(αi, βi) with shape parameters α_i_ and β_i_. After individual *i* making n_j→i_ requests to individual *j* with d_j→i_ interactions that resulted in resource donations, the posterior distribution will follow B(α_i_+d_j→i_, β_i_+n_j→i_). As in any Beta distribution, the expectation made from the perspective of individual *i* is therefore:

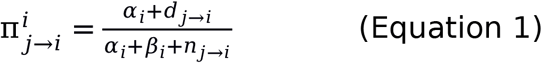

Individual *i* also estimates *j* ‘s belief in *i* ‘s own trustworthiness, π^i^_i→j_ (note that we use here the subscript ‘i→j’). It is calculated as a Bayesian estimate of the probability that individual *i* donates resources when individual *j* has placed a request. This depends on how often individual *i* has donated resources (d_i→j_) when individual *j* asked for it (n_i→j_). The expectation made from the belief of individual *i*is therefore:

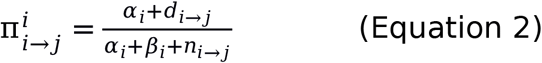

The strength of the same social bond is estimated by individual *j*, **∏**^j^(i,j), in a similar way (but this time with shape parameters α_j_ and β_j_) as the product of:

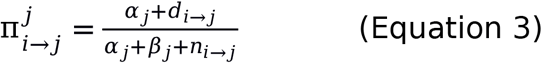

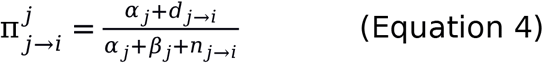

#### Cooperative trait values and social bonds with strangers

In the case where individual *i* encounters a stranger *j*, individual *i* gives the social bond an *a priori* estimate of **∏**^i^(i,j) = π^i^_j→i_ x π^i^_i→j_ = (α_i_ / (α_i_+β_i_))^2^ (because n_j→i_ = n_i→j_ = d_j→i_ = d_i→j_ = 0). Higher values of this prior social bond imply a greater propensity to ask resources from, or to set aside resources for, strangers. The remaining task then is to link this to the traits of individual *i* (note that the trait of *j* does not matter, as there is no information yet on its behaviour).

Trait A_i_ reflects the shape parameters of the prior probability distribution when asking for resources, and G_i_ when giving resources. We also specify a parameter (identical for all individuals) F, which describes the strength of the prior (i.e., high values of F make it more difficult to shift away from prior belief). The bond in the context of asking for resources is formed as α_i_ = A_i_ x F and β_i_ = F (the former switches to α_i_ = G_i_ x F when giving resources). Substituting these values in Equations 1-4 implies that individual *i* uses Ai^2^ (or Gi^2^) as its estimate of the strength of the social bond in encounters with strangers, and can be said to behave according to an initial belief that strangers will help with a probability proportional to A_i_ (or G_i_).

### Processes

At each time step, foraging is followed by resource donations, social bond updating, mortality and reproduction.

#### Foraging

Each time step begins with independent foraging, which ends with each individual in an unsuccessful state with probability P_unsucc_. We assume that foraging success (=1–P_unsucc_) is independent of each individual’s previous foraging successes or previous interactions with group members.

#### Resource donations

If available with probability 1-P_unav_, each successful individual, regardless of her trait A_i_ and G_i_, set aside a total of R resources for donating to others (while consuming the rest of foraged resources individually; we do not model this consumption explicitly, but assume that it allows successful individuals to survive without needing help from others). This total amount R is divided into n_give_ resource ‘packages’ of magnitude r_give_ each (thus R = n_give_ x r_give_). Each of the n_give_ packages are set aside to be donated to a specific individual. The probability that any given individual is assigned by the focal individual to be the intended recipient is proportional to the strength of the social bond as estimated by the focal successful individual (schematized in Fig. 1a).

**Figure 1:**
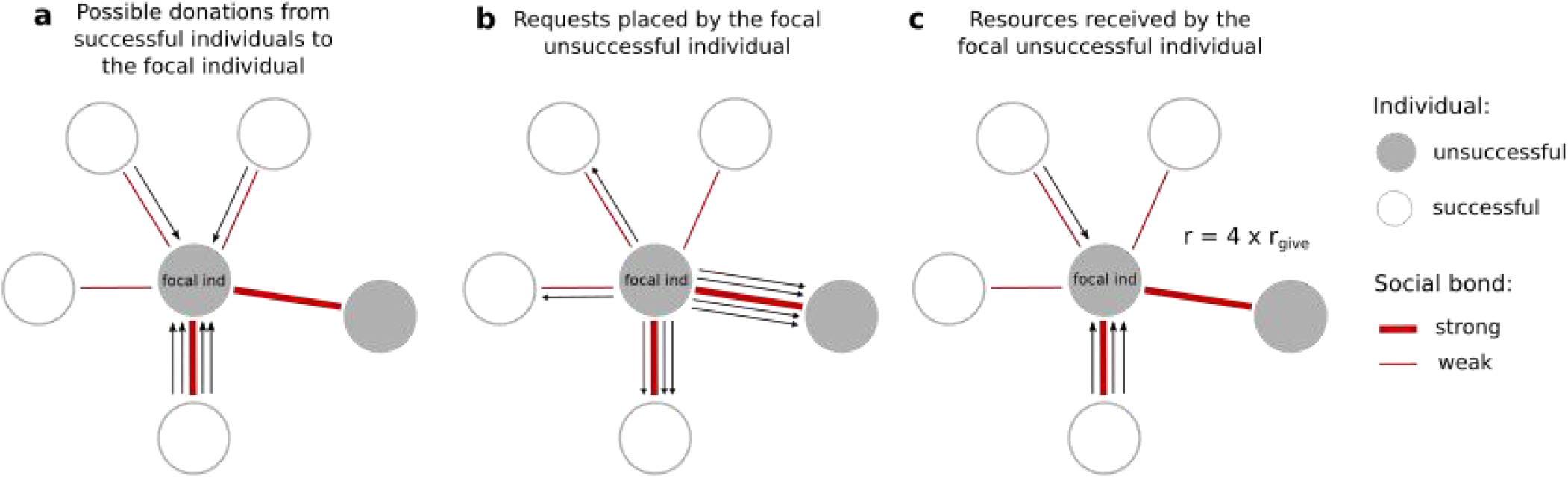
Resource donations from successful individuals to a focal unsuccessful individuals (in the middle of each subfigure). In this example, the focal individual has built strong social bonds with two other individuals, one of them being unsuccessful at this time step. For simplicity, this example assumes that social bonds between each pair of individuals are perceived in the same way by both individuals.

Individuals by definition do not have an interaction history with strangers. Nevertheless, the social bond with a stranger is estimated based on a prior belief (based on G_i_). Depending on the value of G_i_ of the successful individual, the above rules therefore make it possible that one (or more) of the n_give_ packages is set aside to be donated to a particular stranger. Whether an interaction history is existing or not, a focal individual may set aside several packages (or even all n_give_ of them) in the direction of a single recipient, though this is highly unlikely in the case of strangers. Strongly targeted giving is typical for individuals who have already formed a single strong social bond with a specific partner.

Unsuccessful individuals use the social bond network analogously, from their own perspective (schematized in Fig. 1b). Each unsuccessful individual places, during the same time step, a total of n_ask_ requests for help, directed to specific other individuals. Analogously to the above procedure, interactions are preferentially directed towards individuals with whom a strong social bond has been built, but may be directed towards strangers based on a novel social bond estimation that depends on the focal unsuccessful individual’s A_i_ value. Also, analogously to the successful individuals setting aside resources in a targeted fashion, unsuccessful individuals can target several or even all of their n_ask_ requests in the direction of the same individual in the network.

Note that the above decisions of setting aside resources, or placing requests for resources, are done without information on anyone else’s most recent success, or on their decisions to set aside resources or to request them. This means that some of the resources set aside for donation by the successful individuals will not be matched by a request from the intended recipient (who may be successful and thus not needy, or may be unsuccessful but direct requests in some other direction). Likewise, some of the help requests are not matched by willingness to donate, either because there is nothing to donate as both the requester and the target of the request were unsuccessful, or because the target of the help request did not set aside any packages for the requester. If there is a match, all matching resource packages are transferred (schematized in Fig. 1c). Note that the limited number of interaction per time step (controlled by parameters n_give_ and n_ask_) is at the origin of a trade-off between quantity and quality of social relationships.

#### Social bond updating

Resource donations associate with social bond updating. We assume that the total numbers of requests and donations (n_j→i_, n_i→j_, d_j→i_, and d_i→j_ for all pairs of individuals *i* and *j*) are updated among pairs of individuals only if (1) one individual is successful and the other is unsuccessful, and (2) if the successful individual is available and able to give resources. Individuals refine their estimates of the strengths of social bonds with individuals with whom an interaction took place or could have taken place. The total numbers of requests and donations are updated, leading to new estimate values of the strengths of social bonds, **∏**^i^(i,j) and **∏**^j^(i,j), for all individuals *i* and *j*.

The above rules imply that there is no social bond updating when both partners are unsuccessful. Likewise, not being able to give resources (with probability P_unav_) does not harm relationships.

As another source of volatility of the social environment, we assume that any interaction history can be erased with probability P_erase_ leading to a state as if individuals had never interacted with each other. This differs from death in two ways: the two individuals’ other social bonds are kept intact, and they can also begin rebuilding their mutual social bond. This source of volatility is by far the most unrealistic one we implement and may not be found in nature. Nonetheless, we believe it is important to implement it in our model because erasing the history of past interactions is the most parsimonious way to increase social volatility. In particular, this does not change directly age structure (contrary to a direct increase in mortality for instance).

#### Mortality

Mortality has two components. The first, resource-independent component causes each individual, regardless of success and social interactions, to die with probability P_die_. The second, resource-dependent component is only applied to unsuccessful individuals (thus, successful ones always have sufficient resources to survive the second step, regardless of how many resource packages they donated).

Unsuccessful individuals die of starvation during the second round of mortality with a probability that depends on the amount r of resources received during the ‘resource donation’ phase:

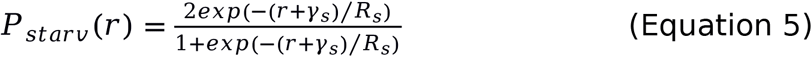

Parameters γ_s_ and R_s_ adjust the risk of death due to starvation. R_s_ scales the overall severity of resource-dependent mortality (R_s_ close to 0 implies that almost all unsuccessful individuals survive; see Fig. 2), while γ_s_ indicates the ‘reserve’ of resources that individuals possess even if they failed both in foraging and in receiving help : if γ_s_ =0, P_starv_(0) = 1. If γ_s_ >0, P_starv_(0) < 1.

**Figure 2:**
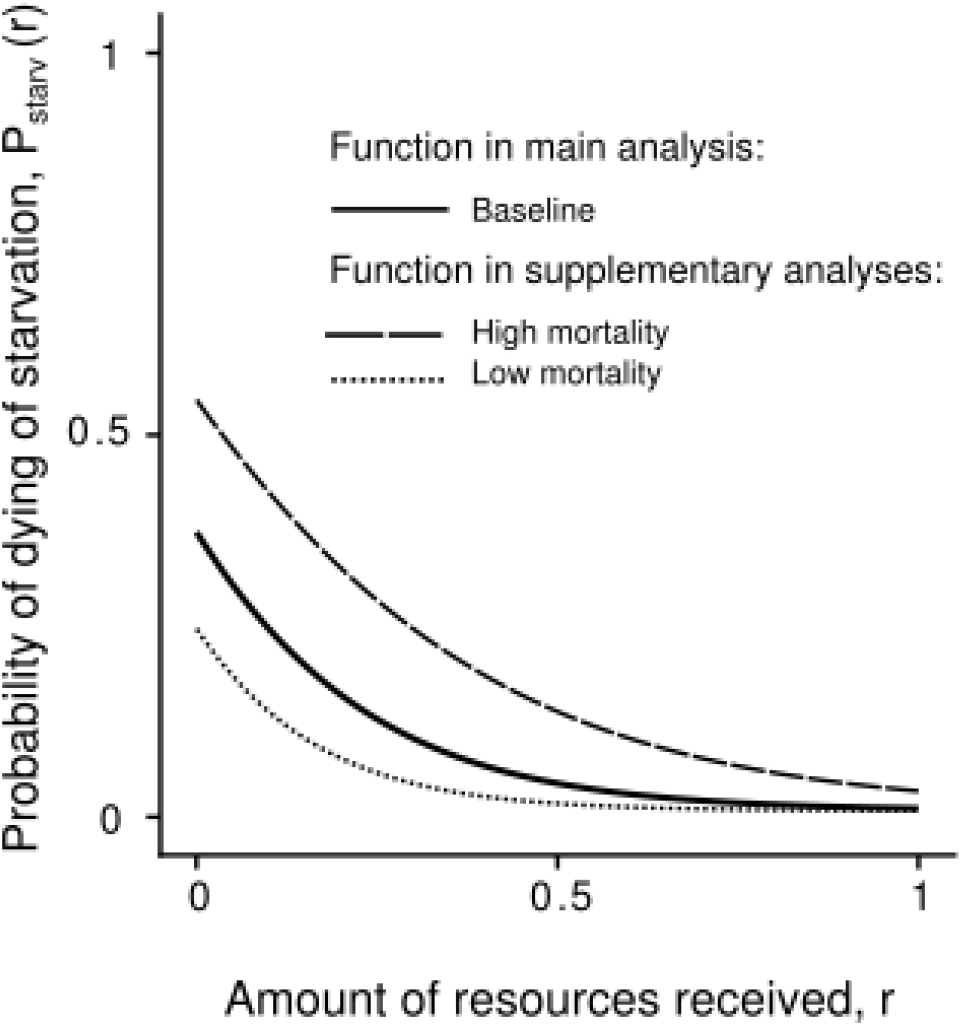
Probability of dying of starvation depending on the amount of resources received by the unsuccessful individual. Baseline function: (γs, Rs)=(0.3,0.2). Other functions implemented in supplementary analyses: (γs, Rs)=(0.3,0.15) (low mortality), and (0.3,0.3) (high mortality).

#### Reproduction

Reproduction is clonal. Each vacancy, created by death, is filled with an offspring, whose mother is a randomly chosen living individual (thus selection acts on survival; there is no differential success among the living potential parents). The offspring traits A_i_ and G_i_ are drawn from truncated normal distributions with means equal to the corresponding parental trait value (reflecting inheritance), with standard deviation σ (reflecting mutation), and with A_i_ and G_i_ constrained to be in the interval [0, 1].

In our main simulations, individuals must build their social network from scratch. In contrast, we assumed in supplementary simulations that newborn individuals initially have strong social bonds with their mother, their sisters or their mother’s partners (social inheritance). Although deviating patterns occur in special cases (that are biologically not likely scenarios, see caption of Fig. S1), as a whole our main message remains robust whether the initial network of a newborn is zero (the network has to be built from scratch) or non-zero (Figs. S1 and S2).

### Simulation experiments

We aim at investigating how the volatility of the social environment affects the evolution of ‘social bet-hedging strategies’, i.e., strategies that diversify cooperative investments. In particular, we test whether changing volatility from low to high favours a shift from a focused to a diversifying approach to social relationships, following the intuition that focusing investments in a single most-profitable partnership is risky if partners often disappear. More precisely, we vary the values of four parameters that determine the volatility of the social environment: the probability P_erase_ of forgetting all information about past interactions with any given individual, the probability P_die_ of dying by chance (i.e. irrespective of foraging success), the probability P_unsucc_ of being unsuccessful, and the probability P_unav_ of being unavailable when successful. The intuitive prediction is confirmed if high values of these parameters lead to the evolution of high trait values A_i_ and G_i_.

We run the model for 5 million time steps. We assume that A_i_=G_i_=0.02 for all individuals i initially (but note that initial variation in trait values does not change qualitatively our results; Fig. S3), and we consider mutations of small effect size (σ = 10^−4^). And, unless stated otherwise, we implement a group size of N=500, a baseline resource-dependent mortality function as shown in Fig. 2, a maximum amount of resources given by each successful individuals R=1, numbers of interactions n_give_=n_ask_=100, and the strength of the prior F=1. A stable social environment is defined by parameters P_erase_=0, P_die_=0, P_unav_=0, and with the only source of mortality determined by cooperative relationships such that P_unsucc_=0.1. Any simulation characterized by higher values of P_erase_, P_die_, P_unav_ and P_unsucc_ is referred to as simulations modeling a volatile social environment. For each combination of parameters tested, we run 30 simulation replicates.

In accordance with the definition of a ‘social bet-hedging strategy’, individuals with higher propensities to interact with strangers (high A_i_ and high G_i_) ultimately diversify cooperative investments across more partners (leading to a higher proportion of individuals with whom the probability of interactions is higher than with a stranger) while building weaker social bonds with each partner (leading to a lower probability to interact with each of those many partners) (Fig. S4).

To describe simulation outputs, we assume that a partnership has been built as soon as an individuals is ten times more likely to interact with this individual rather than with a stranger. The quality of partnership refers to the exact likelihood of interacting with a partner relative to that with a stranger. As noted above, we give the same weight to the perspective of each individual within a pair, and we define the quality of partnership between individuals *i* and *j* as:

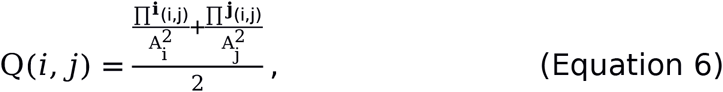

reflecting the degree to which individuals are more likely to interaction with each other than with a stranger. Note that we get qualitatively the same result with **∏**^i^(i,j) (resp. **∏**^j^(i,j)) relative to G_i_^2^ (resp. G_j_^2^). As noted above, we assume that a partnership has been built as soon as Q(*i,j*)>10, with individuals being ten times more likely to interact with each other than with a stranger. This assumption is made for the purpose of describing the outcome of our simulations and does not affect the simulation. In our simulations, low traits values A_i_ and G_i_ leads to more focused cooperative investments than high traits values A_i_ and G_i_, regardless of the definition we use to define partnership establishment (Fig. S4).

## RESULTS

### Evolutionary outcome under stable vs. volatile social environments

In accordance with the definition of a ‘social bet-hedging strategy’, individuals with higher propensities to interact with strangers (high A_i_ and high G_i_) ultimately diversify cooperative investments across more partners while building weaker social bonds with each partner (see examples of emerging social network in Figs. 3 and S5).

**Figure 3:**
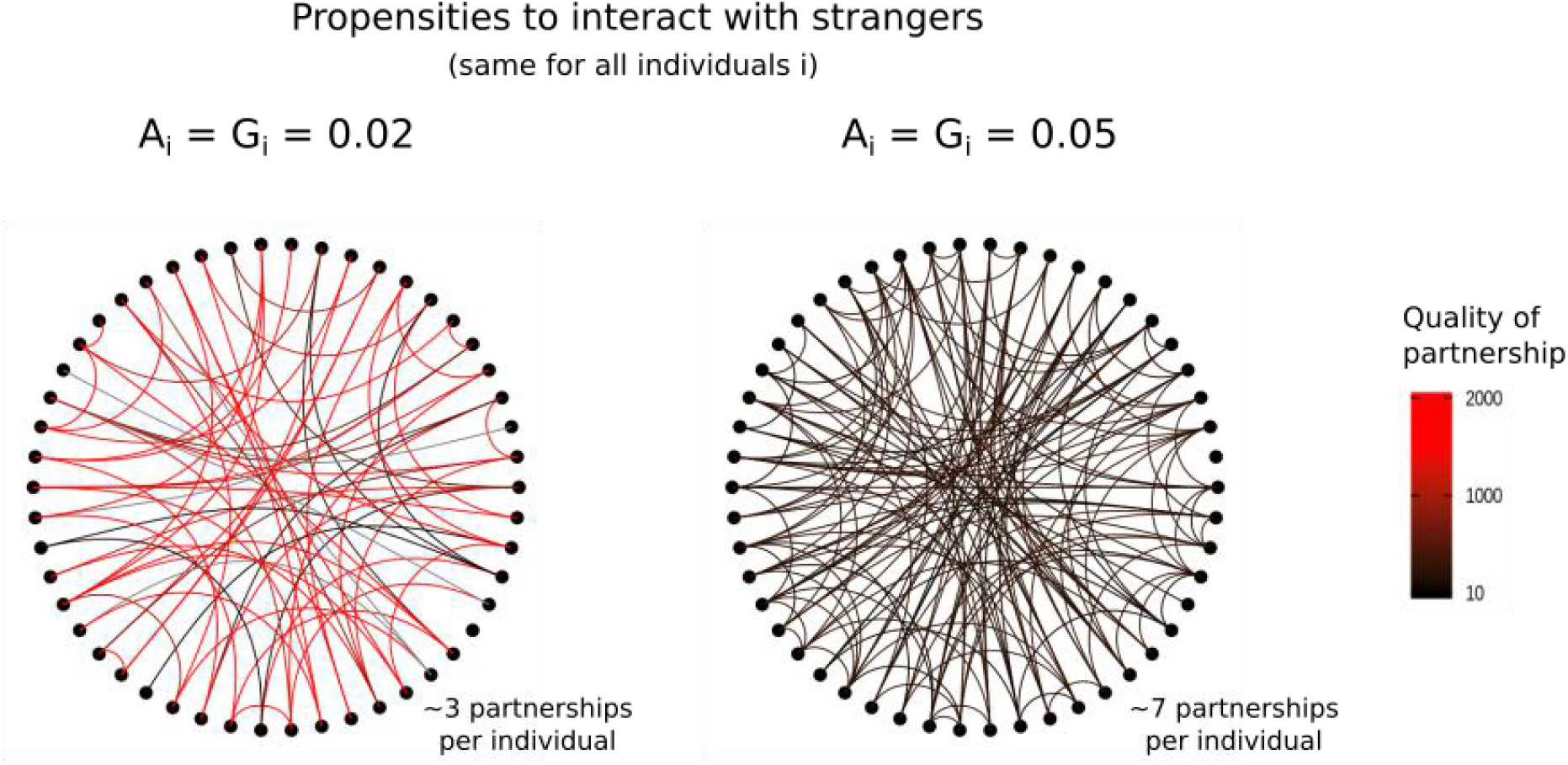
Examples of emerging social networks depending on the socializing behaviours of individuals under a stable social environment (see examples under volatile social environment in Fig. S5). Each link represents a partnership between individuals. The color of the link represents the quality of the partnership -- i.e., Q(i,j) for all pairs of individuals i and j. We assume that a partnership has been built as soon as an individuals is ten times more likely to interact with this individual rather than with a stranger (Q(i,j)>10). Here, N=50.

Contrary to what we expected, a volatile social environment does not favour such diversifying cooperative strategy. Regardless of the type of volatility (P_erase_>0, P_die_>0, P_unsucc_>0.1 and P_unav_>0), higher volatility selects for lower, not higher, values of A_i_ and G_i_ (Fig. 4). In other words, the tendency to diversify cooperative investments is the lowest in the most volatile environments. Our findings (Fig. 4) are not anomalous special cases; we find similar evolutionary outcomes when we change the group size (but note that in a very large group, individuals have to focus investment on few individuals to build partnerships, leading to low trait values A_i_ and G_i_; Fig. S6), the strength of prior expectations (Fig. S7), the maximum number of donations and requests per time step (Fig. S8), the extent of resource-dependent mortality (Fig. S9), and the shape of the resource-dependent mortality function (Fig. S10).

**Figure 4:**
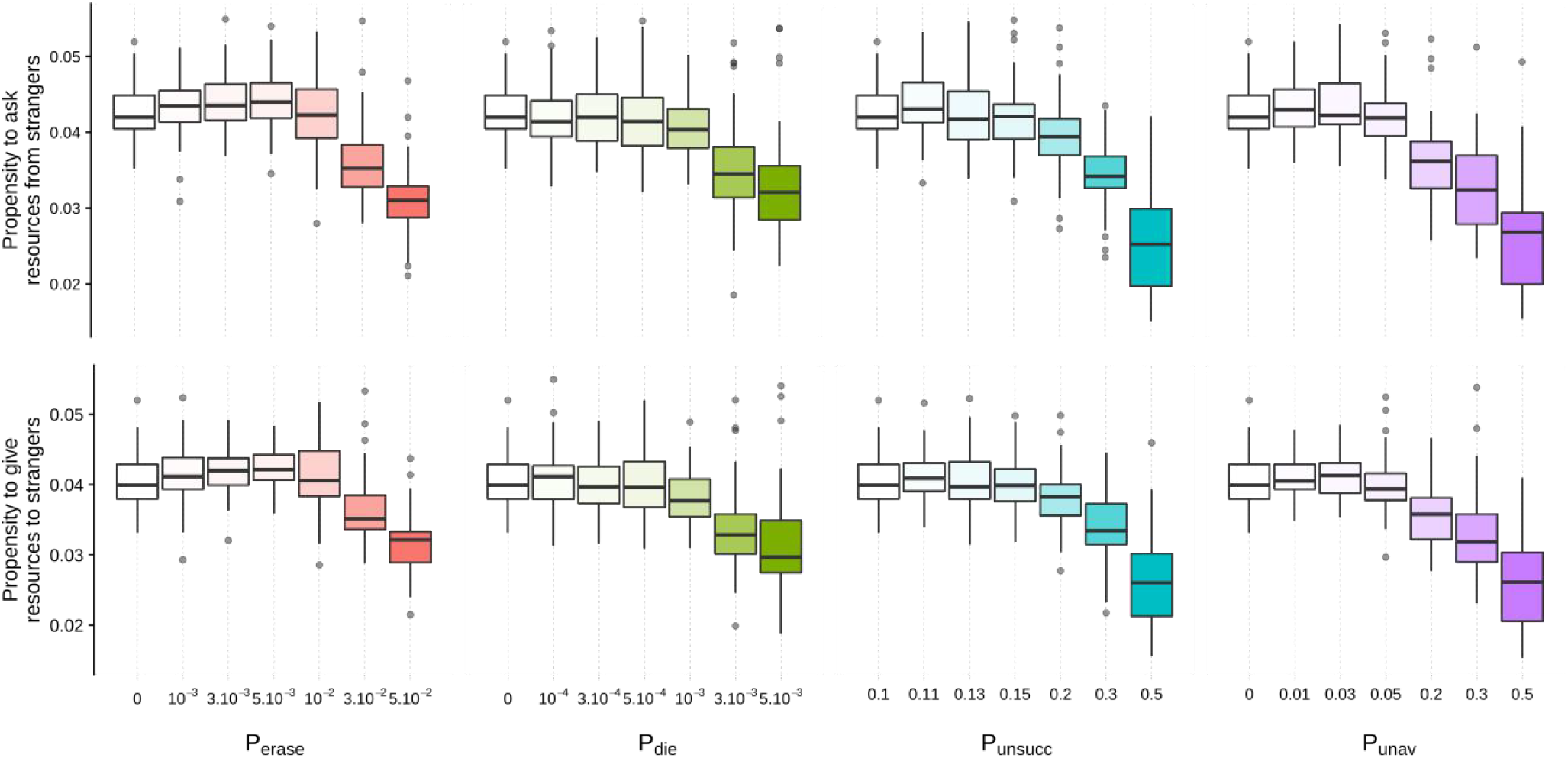
Trait values, A_i_ and G_i_, at evolutionary equilibrium reached after 5 million time steps. High propensities to interact with strangers (high A_i_ and high G_i_) are indicative of a social bet-hedging strategy. For high values of P_erase_, P_die_, P_unsucc_ and P_unav_, the social environment is volatile. Under these conditions, the evolutionary equilibrium is characterized by cooperative strategies focusing on the quality of each partnership rather than on the quantity of partnerships (low Ai and low G_i_, i.e. the opposite of social bet-hedging).

We performed additional simulations where newborn individuals have a strong social bond with their mother, their sisters, or their mother’s partners (Supp. Figs. S1 and S2) and do not build their social network from scratch. In most cases, volatile social environments keep favoring more focused cooperative strategy than do more stable social environment (at the exception of increased P_erase_, which is the most unrealistic form of volatile social environments, as detailed in Fig. S1). Interestingly, we show that a focused cooperative strategy translate into a higher proportion of partnership between relative vs. unrelative individuals when partnership between kins occurs, but into a lower proportion of partnership between relative vs. unrelative individuals when social inheritance takes place (Fig. S2).

### Demographic feedback under a volatile social environment

It is clearly necessary to explain why a volatile social environment leads to the evolution of a strategy focusing cooperative investment on few partners. We illustrate this with an example, where we trace the life of a single mutant individual *i* with deviating traits A_i_, G_i_ when all other population members *j* have trait values A_j_=0.035 and G_j_=0.035. For the mutant *i* we consider traits values A_i_, G_i_ ϵ {0.02, 0.035, 0.05}. We then assess the characteristics of the social relationships and the survivorship of this mutant as it ages. Age is defined based on time steps at unsuccessful state; note that this quantification of age is strongly correlated with the true age of the individual based on all time steps (Fig. S11).

Individuals with high A_i_ and G_i_ values ultimately have many partnerships of poor quality (first and second rows in Fig. 5). Nonetheless, at early age, these individuals have fewer partnerships than individuals focusing on few partnerships (with low A_i_ and G_i_ values). Focusing on few partnerships speeds up social bonding and increases the exploitation of the benefits associated with the existing partnerships, ultimately increasing survival at early age (third and fourth rows in Fig. 5). Although the focused approach carries some risk, they are smaller than the reduction in early performance if attempting to diversify when one’s own network is still in the first stages of being built. This occurs even if newborn individuals do not need to build their social network from scratch; exploiting the benefits associated with the pre-existing social bonds (e.g., with the mother; Fig. S12) at the expense of diversifying cooperative investments increases survival at early age.

**Figure 5:**
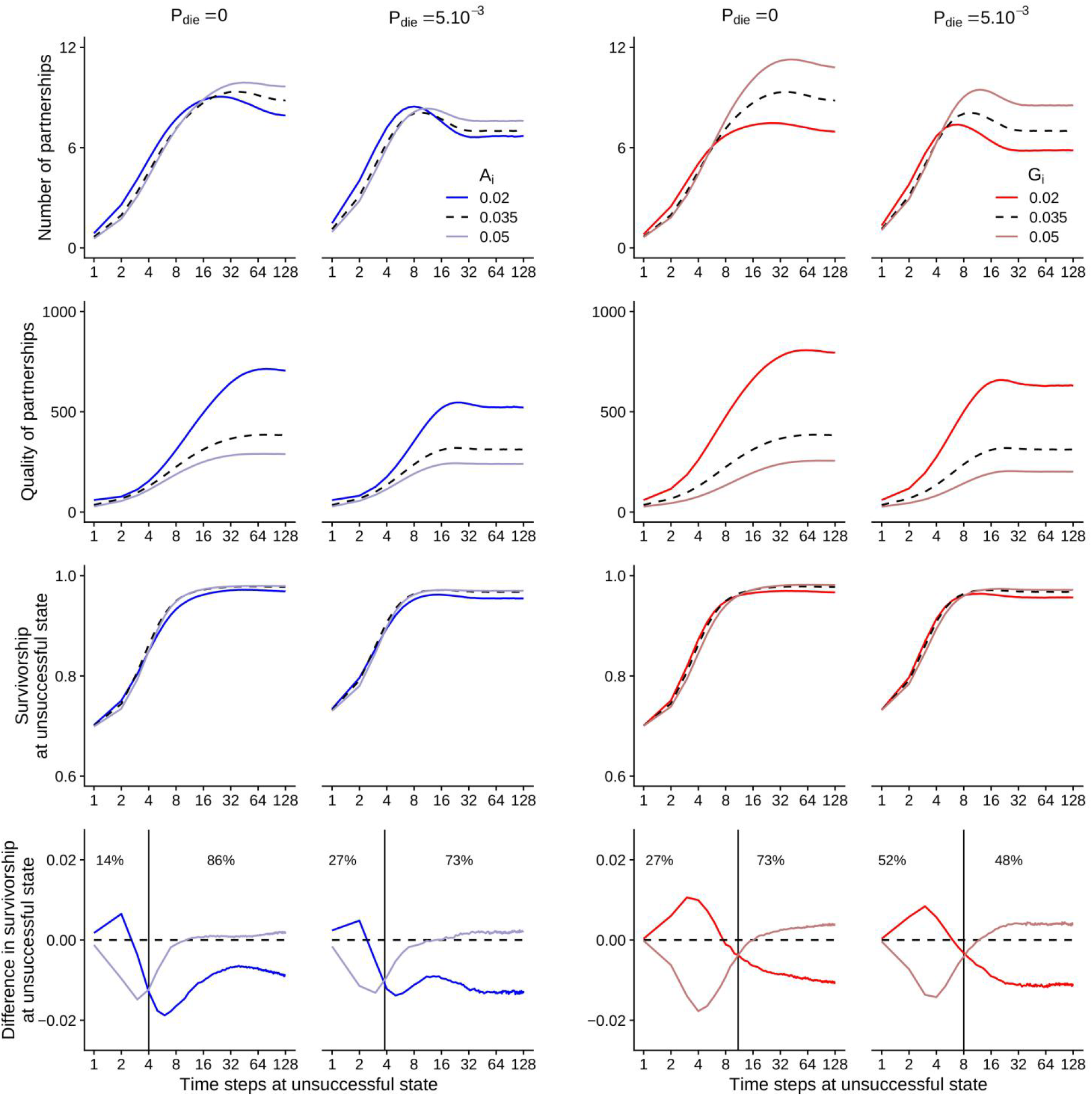
Characteristics of social relationships and survivorship of one individual *i* depending on its age and on its trait value in a population where A_j_ = G_j_ = 0.035 for all other individuals *j* either under a stable social environment or under a volatile social environment (P_die_ = 5.10^−3^). The mutant individual *i* differ either by its trait value A_i_ (in blue) or by its trait value G_i_ (in red). From top to bottom, we represent the quantity of relationships (following the criterion Q(i,j)>10), the quality of relationships (Q(i,j)), survivorship, and difference of survivorship compared with other individuals in the population. In the last row, the vertical line delimits age categories where investing in few relationships (trait value = 0.02; dark colour) is beneficial compared with investing in many relationships (trait value = 0.05; light colour) and vice versa. Percentages correspond to the proportion of individuals in each of those two age categories.

In other words, focusing on few partnerships appears necessary at early age, whereas building many partnerships proves beneficial at older ages once the many partnerships have been built.

Under a volatile social environment (when P_die_ = 5.10^−3^ in Fig. 5; but this is also true for high P_erase_, P_unsucc_ and P_unav_ as shown in Figs. S13, S14 and S15), diversifying cooperative investment benefits old but not young individuals (old individuals with high A_i_ and G_i_ trait values have a higher survivorship than old individuals with low A_i_ and G_i_ trait; last row in Fig. 5). In that sense, the premises on the benefits associated with a ‘social bet-hedging strategy’ hold: in volatile social environments such diversifying cooperative strategy proves beneficial late in life because is reduces the temporal variance in cooperative returns caused by unpredictable changes in partner availability (Carter et al. 2017). Nevertheless, the evolution of such a cooperative strategy does not depend only on the performance of individuals belonging to a certain age group. Here, a key question is how much of one’s life one will spend being ‘young’ versus ‘old’. As a whole, the selection gradient changes as the age distribution changes (Fig. S16).

Overall, high mortality increases the proportion of young individuals relative to old individuals. As a result, under a volatile social environment, fewer individuals reach old ages where they can reap the benefits of diversifying cooperative investment. The details differ between settings: a volatile social environment may associate with high mortality directly (high P_die_), indirectly (high P_erase_ and high P_unav_) or both directly and indirectly (high P_unsucc_; via resource-dependent mortality and via a lower pool of helpers). Regardless of the specific route to short lives, volatility means that relatively few individuals reach sufficiently old age to reap the benefits of a diversified approach to network building. Instead, speeding up the socializing process and exploiting existing partnerships at an early age matters the most (see percentage in last row of Fig. 5), even if this proves deleterious later in life (should the individual still be alive, which is relatively unlikely under high volatility). This explains why a ‘focused’ rather than diversifying approach to cooperative investment is beneficial under a volatile social environment.

## DISCUSSION

Our model shows the fragility of the intuitive prediction that stable social environments should favour individuals that develop few but strong relationships, while volatile social environments favour those whose social networks end up ‘shallow but broad’. Our model shows that the premises on the benefits associated with a diversifying cooperative strategy hold: ‘shallow and broad’ social networks can prove beneficial because they reduce the temporal variance in cooperative returns caused by unpredictable changes in partner availability (as argued by Carter et al. 2017). Nevertheless, our model also highlights that the evolution of cooperative strategies does not depend solely on the performance of the social networks once they are built. When accounting for the costs associated with the build-up of such social networks, selection can lead to the precise opposite outcome: volatility selects for focused network building.

The reason is clear once reciprocity theory is linked to theories of senescence, specifically, antagonistic pleiotropy (Williams 1957, Flatt & Partridge 2018). Diversifying cooperative investments can be beneficial late in life (once the individual has built its social network), but this is preceded by a substantial cost early in life while the network has to be built. Since the probability to reach a sufficiently old age is low in volatile environments, the negative effect early in life predominates, and the successful strategy is one that focuses cooperative investments on few partners. Focusing allows individuals to build strong partnerships more quickly while also exploiting existing social bonds (including those with relatives, and inherited ones). It is notable that a strategy that focuses reciprocal interactions on few individuals is able to spread in a population, even though it clearly has the potential to lead to a disastrous loss of all ‘friends’ for some individuals. Our model accounts for this cost, and simply shows that the beneficial effects of a focused strategy are on average better than those of a diversifying one.

Our results highlight the importance of population age structure for the evolution of pleiotropic traits that are beneficial early in life but deleterious later. This concept features strongly in senescence research, where ‘antagonistic pleiotropy’ hypothesizes that alleles that enhance fitness early in life but are detrimental later can be favoured because selection is stronger early in life than late in life (Williams 1957). An increase in extrinsic sources of mortality makes it less likely that an individual will survive from birth to an old age, increasing selection favouring those alleles at pleiotropic genes and favoring more rapid senescence (Williams & Day 2003). Our model highlights that a similar evolutionary force occurs under a volatile social environment when cooperative strategies are pleiotropic, analogous to recent developments in life history theory where processes that are optimized for early-life function can become detrimental later (Maklakov & Chapman 2019).

Pleiotropic effects are also discussed in the field of social behaviour, where it has been suggested as a mechanism stabilizing cooperation in slime moulds and bacteria (Foster et al. 2004, Dandekar et al. 2012; but see Dos Santos et al. 2018). These cases, however, do not have the same pleiotropic structure as the one we consider. We focus on a situation where pleiotropy is clearly age-dependent, and our question is also different: we do not consider whether cheats can spread and destroy cooperation, instead we ask how reciprocal cooperation deals with the quantity-quality tradeoff ‘beyond the dyad’ (as advocated by Connor 2010). It is of interest to note that the field of social behaviour is generally starting to realize that the social environment is very likely different for individuals differing in their age; Croft et al. 2021 discuss this with respect to kinship (see also Rodrigues 2018). Our work shows that the social state, i.e. the position of an individual within its network of social partners and the properties of this network, can be age-dependent in a manner that can switch selection from favouring narrowing this network down or broadening it further.

In our model, pleiotropy arises from differences in the exploitation of existing social bonds but also from differences in the speed at which social bonding takes place. What is known as ‘social bet-hedging strategy’ refers to a diversifying approach to social relationships, and via the quality-quantity trade-off, we have shown it associates with slow social bonding. Cooperation typically relies on some form of assortment (Fletcher & Doebeli 2009, Marshall 2011, Garcia et al. 2015), which involves individual recognition and social bond formation (at least in the type of organisms that our model is inspired from). We know very little on how social bonds initially form, especially when they entail investments of time and energy. Social bonding may be characterized by a raise in cooperative investments over time, as long as reciprocal cooperation takes place (Roberts and Sherratt 1998, Fruteau et al. 2011, Carter et al. 2020). In our knowledge, however, the speed at which social bonding takes place has rarely been assessed; in vampire bats, social bonding taking the form of reciprocal grooming and reciprocal donations can take up to 200 days (Carter et al. 2020). Our model shows that the speed at which social bonding takes place is likely to have a strong impact on the evolution of cooperative strategies because most of social bondings occur early in life.

It is interesting to reflect on the pitfalls of intuition. Intuition often involves ‘putting oneself in another organism’s shoes’ — in sometimes fallible ways (Kokko 2017). In the current context, intuition may be based on imagining what one should ideally have done, given the sudden death of a social partner. Clearly, having built a broad network helps to recover future fitness prospects, mitigating the current loss. Strangely, intuition does not prompt us to reflect equally much on the possibility that death might target ‘oneself’ (the focal individual). Yet volatility obviously also strikes, with some regularity, this way, and now the hope is that one did whatever one could to maximize performance until that age; death made performance at later ages unmeasurable and irrelevant. If intuition only considers actions and their benefits among those who keep avoiding death themselves, it falls victim to the well-known effects of ‘survivorship bias’.

Survivorship bias also is a tough problem for the empirical aspects of the question, as data collection on behavioural details can, logically, only be based on observations involving current survivors. While frustrating, this also helps to understand apparent discrepancies between data and our model. Losses of individuals whose networks did not help them avoid death create a process of selective disappearance within each cohort. This makes it exceedingly hard to collect unbiased data: unless one traces social bonds longitudinally and records every death, any analysis among living individuals will pay disproportionate attention to the successful subset of individuals who are presently observable by virtue of being alive — a problem which applies to a wide range of taxa (beyond bats), whenever aiming to document effects of the quantity vs. quality of social relationships (Sutcliffe & Crabbe 1963, Amato 1993, Entwisle et al. 2007, Sato & Zenou 2015, Vacca 2019, Silk et al. 1999, Carter et al. 2017, Ellis et al. 2019, Morrison et al. 2020, Menz et al. 2020, Bond et al. 2021). Whenever disappearance is selective (not random), the problem is exacerbated by the fact that situations where selection is at its strongest also produce the most severe data collection biases. To help solve this conundrum, future empirical studies paying particular attention to the longitudinal aspect of individual lives (and the age-dependent dynamics of their networks) could shed new light onto the question of well performing networks and their temporal trade-offs.

One notable study with a temporal aspect is the one by Testard et al. (2021), where the authors showed that macaques diversified their social relationships after their population was devastated by a hurricane. We believe that the discrepancies between this result and our theoretical predictions can be explained. First, such a dramatic event is far from the level of instability we modelled, and second, the observed changes in cooperative behaviours in those macaques were an example of plasticity, not an evolutionary response to permanent volatility that would select for a different type of behaviour due to individuals routinely dying young. Testard et al’s study therefore highlights some limits of our modelling approach. It is a clear avenue for future work to consider plasticity, as expression of behaviours is often remarkably sensitive to environmental conditions (Snell-Rood 2013).

Plasticity could also make traits age-dependent and also perhaps dependent on the state of one’s own network: cooperative behaviours could change through life. If one’s social network is already broad enough, one may change its cooperative strategy and stop expanding it further. Such age-dependent and/or network state-dependent plasticity has received some empirical support in rhesus macaques, where older females engage less in the social environment compared to younger ones (Brent et al. 2017), and should therefore be investigated in future theoretical studies. Based on our predictions, a strategy focusing cooperative investment on few partners at early age and diversifying cooperative investment on many partners at old age could conceivably be optimal. However, note that a social network of an individual at any point in its life is an accumulation of an entire ‘career’ of work towards it. Even if behavioural changes are possible, it is not clear that an adjustment schedule is able to choose performance in early life such that two goals are simultaneously optimized: to have as good as possible fitness should death happen at a relatively young age, and, should early death not occur, to ‘prepare’ the individual’s social network to allow best possible capitalization of the gains that follow from a switch in strategy (since any attempt to broaden the network must start from where early-age efforts ended). Since it is never clear how long an individual life will last, antagonistic pleiotropy may be unavoidable even under plastic social traits.

While we showed that the volatility of the social environment has little effect on cooperative strategies in very large groups (over 2,000 individuals), this is likely not to be the case when groups are organized into subgroups that merge or break up over time. Such subgroup dynamics is a common feature of society of cooperative animals (e.g., in vampire bats; see a review in Aureli et al. 2008) and may conceivably determine the nature of cooperative strategies at evolutionary equilibrium. In particular, the extent to which subgroup dynamics interact affect age structure and age-dependent effects of cooperative strategies on fitness remains to be formally investigated.

As a whole, our model highlights the importance of population age structure for the evolution of social traits such as cooperative behaviours. The evolution of social traits has been traditionally studied using evolutionary game theory (Maynard Smith 1982, McNamara & Leimar 2020) and quantitative genetics (Wolf et al. 1999, McGlothlin et al. 2014, McGlothlin et al. 2021), without much emphasis on the implication of demography. Recent studies have started to uncover the role of spatial structure for the evolution of social traits (e.g., Débarre et al. 2014, Peña et al. 2016, Su et al. 2019), but little is known on the role of age structure (but see Rodrigues 2018 for a a age-dependent kin-selection model). We appreciate that our individual-based modelling approach comes with its limits, including the abscence of analytical insights; we hope however that the predictions of our model will stimulate further theoretical and empirical investigations assessing the role of population age structure for the evolution of social traits.

## ACKNOWLEDGMENTS

We thank G.G. Carter for discussion and inspiration. We also thank D.R. Farine, M. Servedio, B. Lerch, K. Xu and two anonymous reviewers for comments that have helped improve our manuscript. Simulations were performed on the computing cluster of the Science Cloud platform of the University of Zurich.

