## Supplementary Figures for "Volatile social environments can favour investments in quality over quantity of social relationships"

**Figure S1 (next page):** Trait values, $A_i$ and $G_i$, at evolutionary equilibrium reached after 5 million time steps, assuming newborn individuals do not build social networks from scratch. In all panels, we assume that newborn individuals have initially strong social bonds with their mother (equivalent to social bonds acquired following 100 resource donations out of 100 requests). In panel **b**, we assume that newborn individuals also have strong social bonds with their siblings (equivalent to those acquired following 10 resource donations out of 10 requests), whereas in panel **c**, we assume that newborn individuals inherit their mother's social network to some extent (initial social bonds are computed as if newborn individuals have already placed 10 requests to each individual, and that donations occurred following the appreciation of the mother by each individual). With initial social bonds with relatives, individuals have to focus investment on few individuals to strengthen partnerships with their kins, leading to low trait values $A_i$ and $G_i$ at evolutionary equilibrium. Overall, however, we get the same outcome than without initial social bonds; for volatile social environments, the evolutionary equilibrium is characterized by cooperative strategies focusing on the quality of each partnership rather than on the quantity of partnerships as in the main analysis. For high values of $P_{erase}$, we get a qualitatively different outcome when individuals share the same social network as their mother: a high $P_{erase}$ value leads to a more diversifying strategy than under a stable social environment (**a**, **b**). In that case, when mother and offspring forget all information about their past interactions with each other, they get in strong competition for access to resources from shared partners (without being able to benefit from a strong mother-offspring bond anymore); as a result, a diversifying cooperative strategy proves more beneficial then under a stable social environment. Indeed, when social bonds between mothers and offspring cannot be erased (simulation assuming that 'any bond except mother-offspring' can be erased, in panels **a** and **b**), we get back our main outcome. Given that the scenario with individuals forgeting all information about past interactions is the more unrealistic one and may not describe social network volatility in many empirical system, we can conclude that our main result mostly hold when assuming that newborn individuals do not build social networks from scratch.

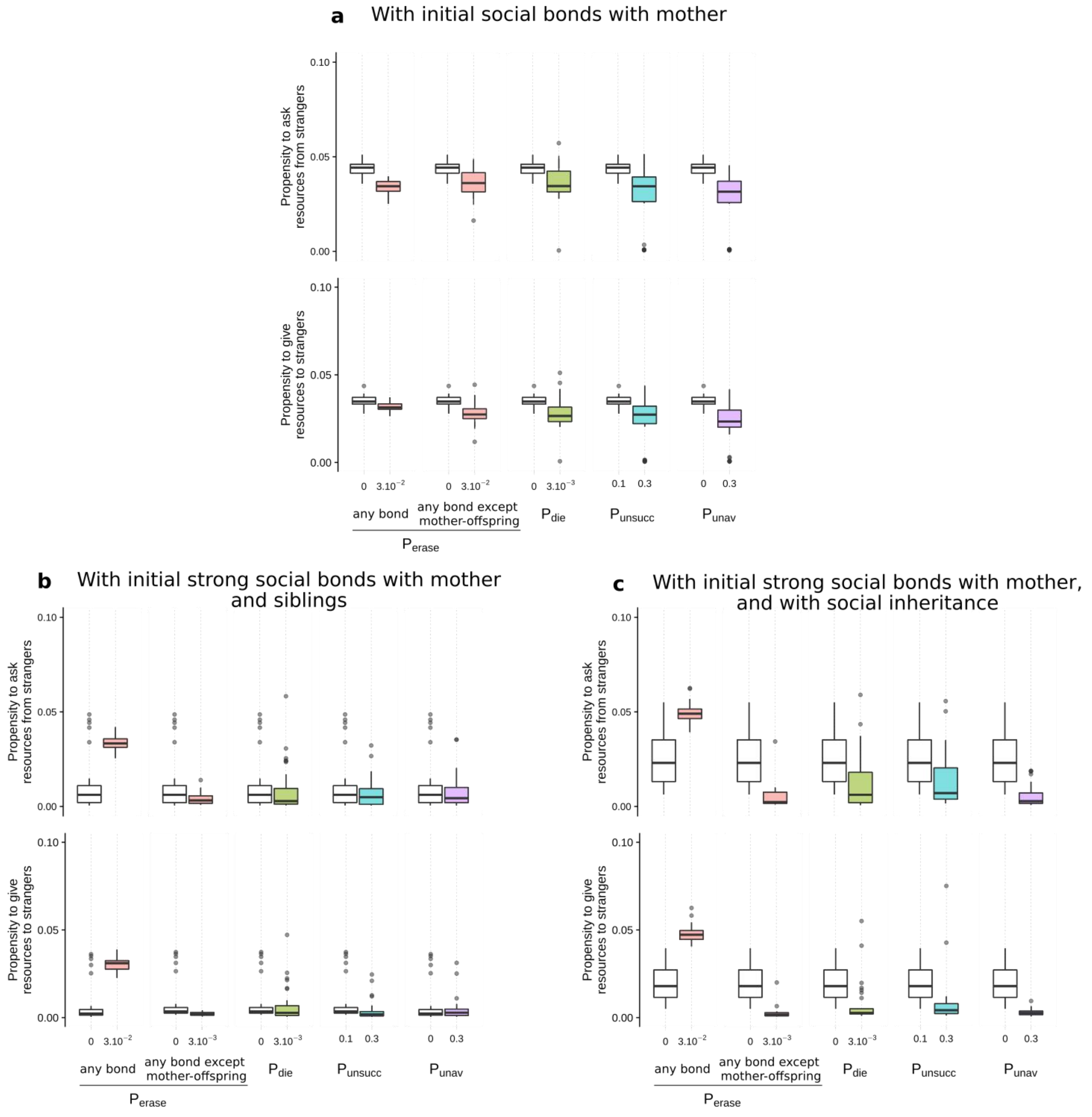
a With initial social bonds with mother
b With initial strong social bonds with mother and siblings
c With initial strong social bonds with mother, and with social inheritance
Propensity to ask resources from strangers
Propensity to give resources to strangers
0.10
0.05
0.00
0
3.10−2
3.10−3
0.1
0.3
any bond
any bond except mother-offspring
Pdie
Punsucc
Punav
Perase

**Figure S2:** Mean proportion of partners that are unrelated at evolutionary equilibrium reached after 5 million time steps, assuming newborn individuals do not build social networks from scratch (in the same conditions as in **Fig. S1**). Interestingly the proportion of unrelated partners is not solely determined by mean trait values, $A_i$ and $G_i$. In unstable environments, low propensities to interact with unknown individuals (low $A_i$ and $G_i$) associate with low proportion of parnerships among unrelated individuals when social inheritance occurs (**a**, **b**). When social inheritance occurs, however, low propensities to interact with unknown individuals actually increases the proportion of partnerships between unrelated individuals (**c**). See Fig. S1 for more details.

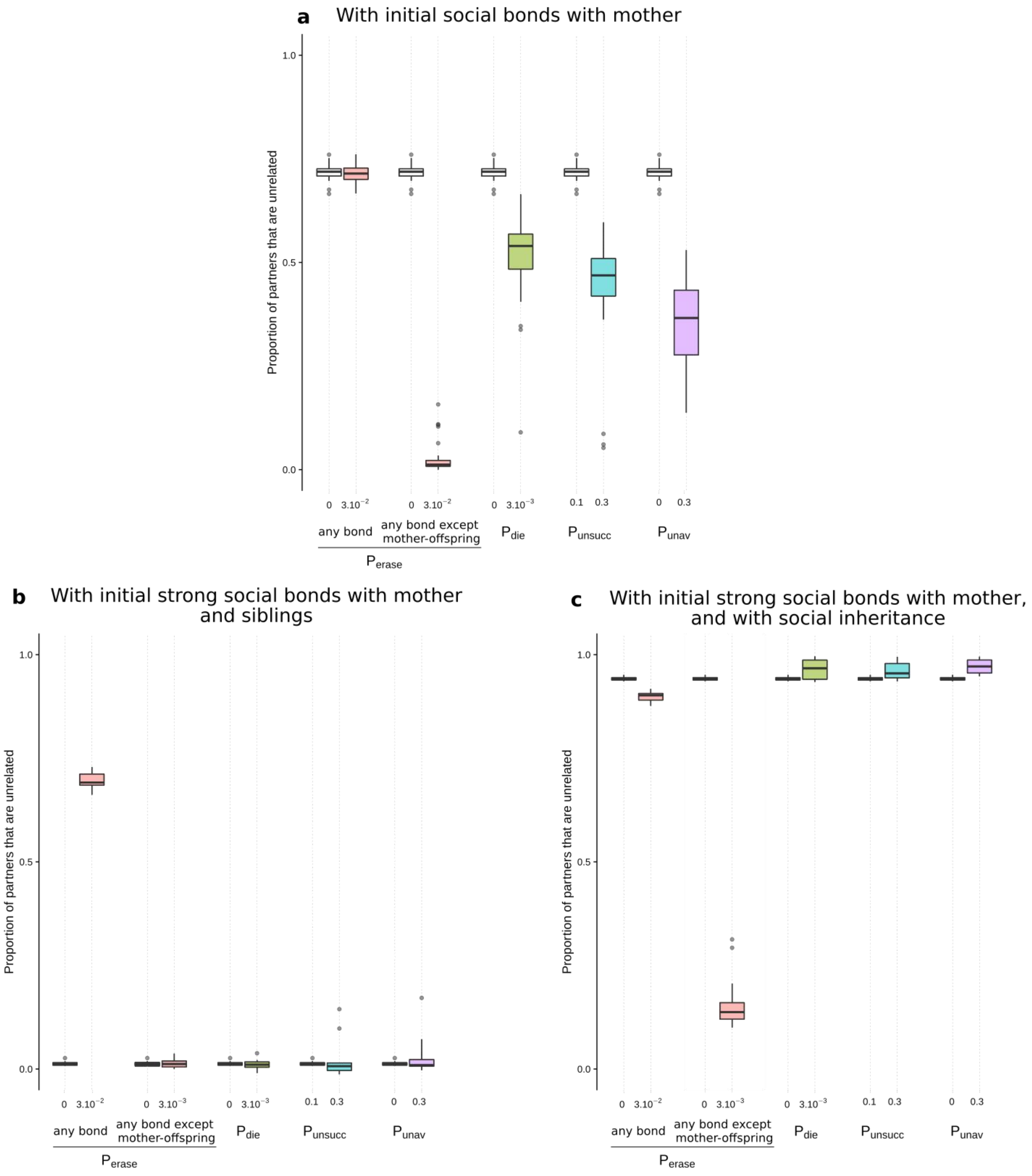

**Figure S3:** Trait values, $A_i$ and $G_i$, at evolutionary equilibrium reached after 5 million time steps, assuming initial variation of the traits between individuals. Instead of assuming that $A_i$=$G_i$=0.02 for all individuals i initially, traits $A_i$ and $G_i$ are initially drawn from truncated normal distributions with means equal to 0.02, with standard deviation equal to 0.01 or 0.05, and with $A_i$ and $G_i$ constrained to be in the interval [0, 1]. In each graph, we represent the initial distribution of trait values in gray. We get the same outcome than without initial variation of trait values; for high values of $P_{erase}$, $P_{die}$, $P_{unsucc}$ and $P_{unav}$, the evolutionary equilibrium is characterized by cooperative strategies focusing on the quality of each partnership rather than on the quantity of partnerships (low $A_i$ and low $G_i$) as in the main analysis.

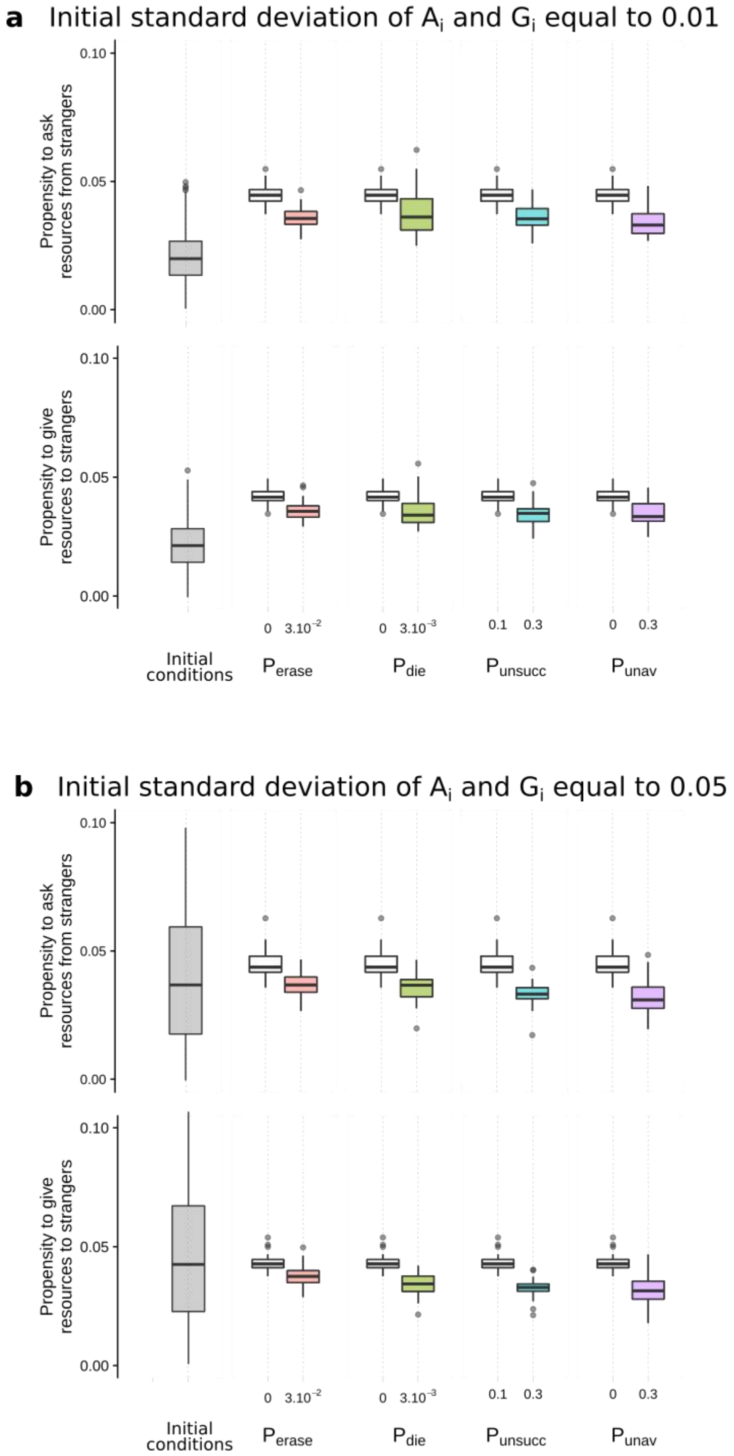

**Figure S4:** Distribution of the likelihood of interacting with individuals relative to that with a stranger, at evolutionary equilibrium reached after 5 million time steps, depending on the socializing behaviours of individuals (propensities to interact with strangers) and on the volatility of the social environment. In each graph, the vertical dashed line represents the arbitrary threshold value that we used to define a partnership: we assumed a partnership has been built as soon as an individuals is ten times more likely to interact with this individual rather than with a stranger. In accordance with the definition of a 'social bet-hedging strategy', individuals with higher propensities to interact with strangers (high $A_i$ and high $G_i$) ultimately diversify cooperative investments across more partners (higher proportion of likelihood of interactions greater than one) while building weaker social bonds with each partner (lower magnitude of likelihood of interactions). This outcome is also captured using our definition of partnership establishment; high $A_i$ and high $G_i$ result in social networks with more partnerships of low quality (see text within each graph). Additionally, under a volatile social environment, individual lifespan is short, and the proportion of likelihood of interations greater than one (and therefore the number of partnerships) remains very low because individuals do not have time to build many relationships. Here, N=500.

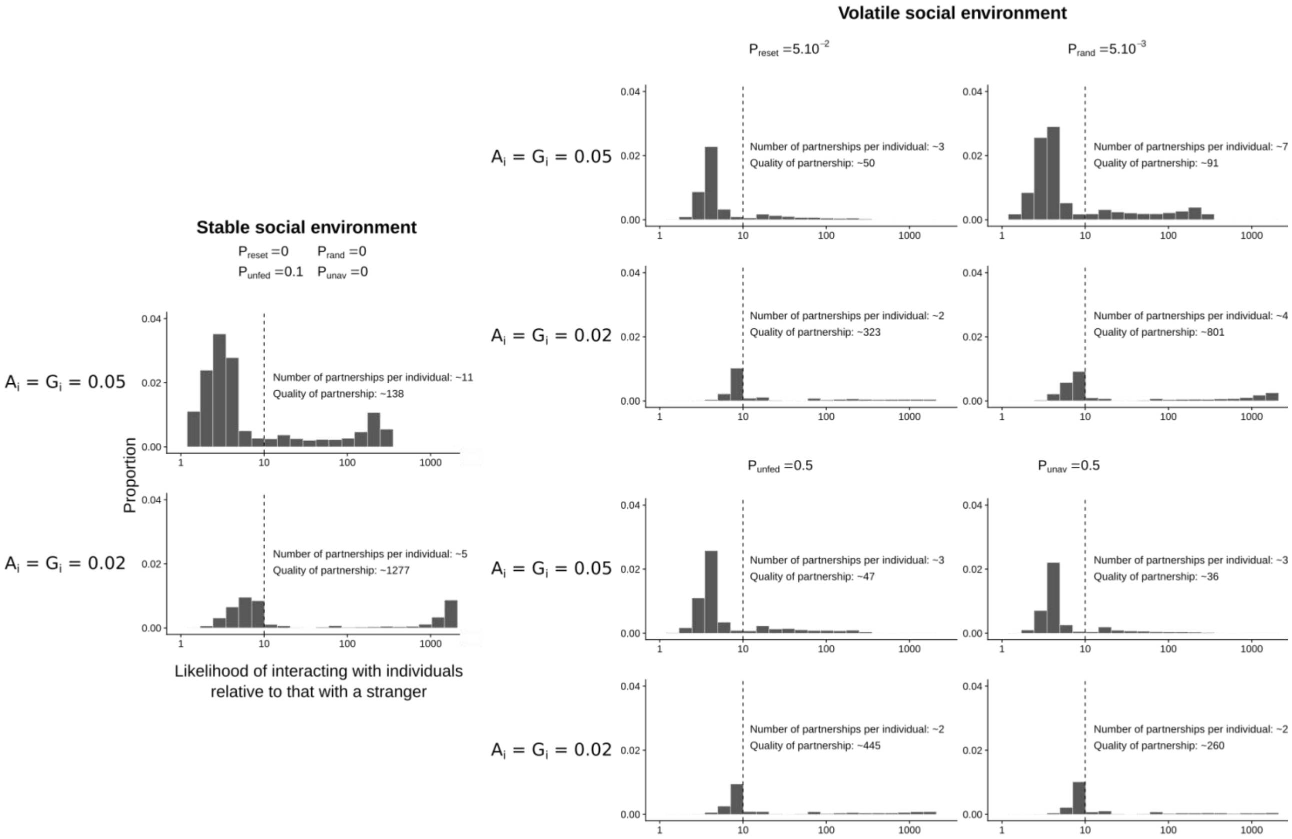

**Figure S5:** Examples of emerging social networks depending on the socializing behaviours of individuals (propensities to interact with strangers) and on the volatility of the social environment. See caption of Figure 3 for more details. Under a volatile social environment, individual lifespan is short, and the number of partnerships remains very low because individuals do not have time to build many relationships. Here, N=50.

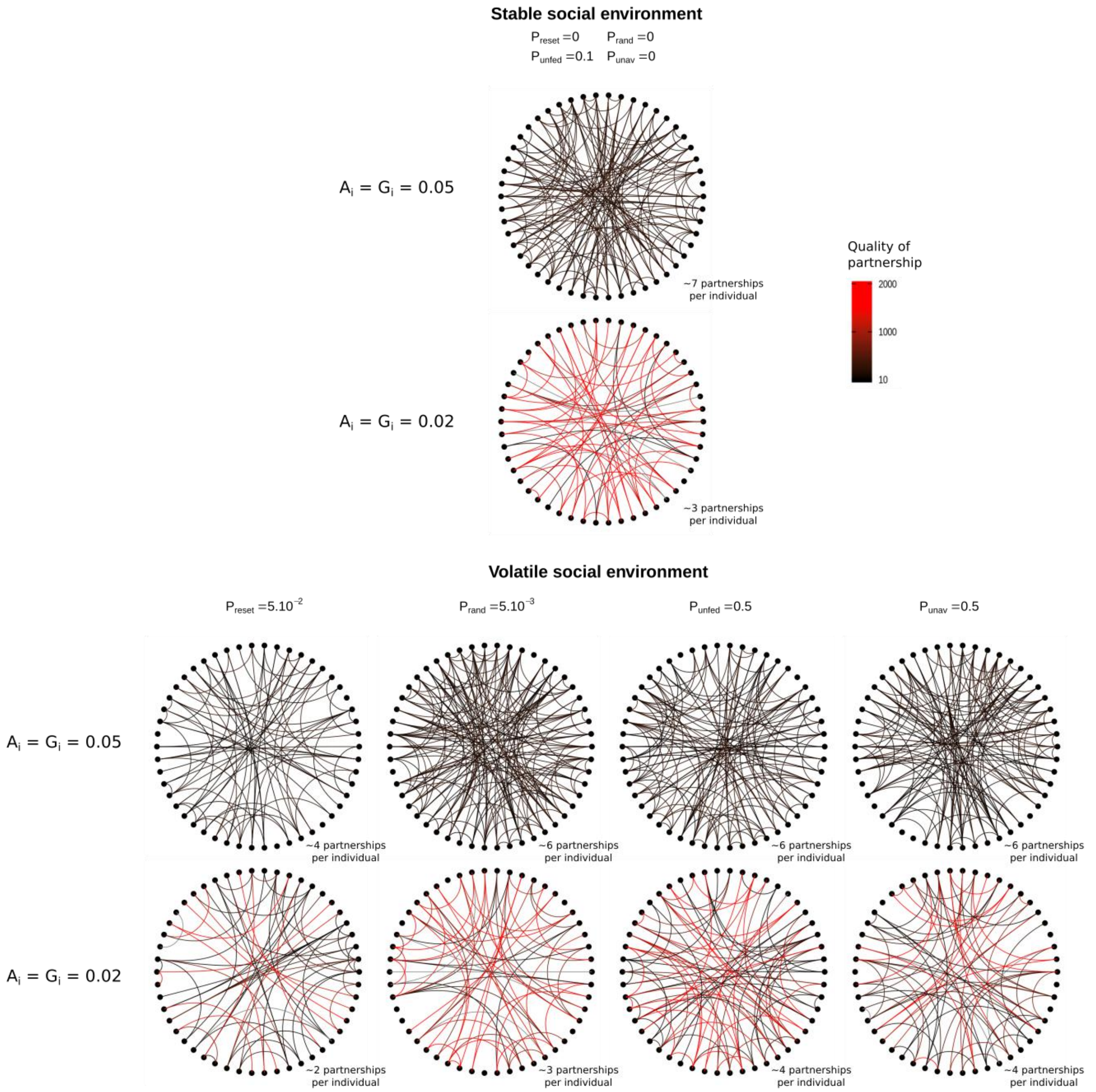

**Figure S6:** Trait values, $A_i$ and $G_i$, at evolutionary equilibrium reached after 5 million time steps, for different group sizes. In a large group, individuals have to focus investment on few individuals to build partnerships, leading to low trait values $A_i$ and $G_i$, because the risk of diluting cooperative investments is too high. For high values of $P_{erase}$, $P_{die}$, $P_{unsucc}$ and $P_{unav}$, the evolutionary equilibrium is characterized by cooperative strategies focusing on the quality of each partnership rather than on the quantity of partnerships (low $A_i$ and low $G_i$) as in the main analysis. For very large groups, however, the volatility of the social environment has little effect on the traits values $A_i$ and $G_i$ (that tend toward 0) at evolutionary equilibrium (for N = 5,000 here). Remember that in our model, individuals may interact with all individuals of the group; in nature, such large group are often divided in subgroups, whose dynamics (fusion and fission) may affect the evolution of cooperative strategies (not explored here).

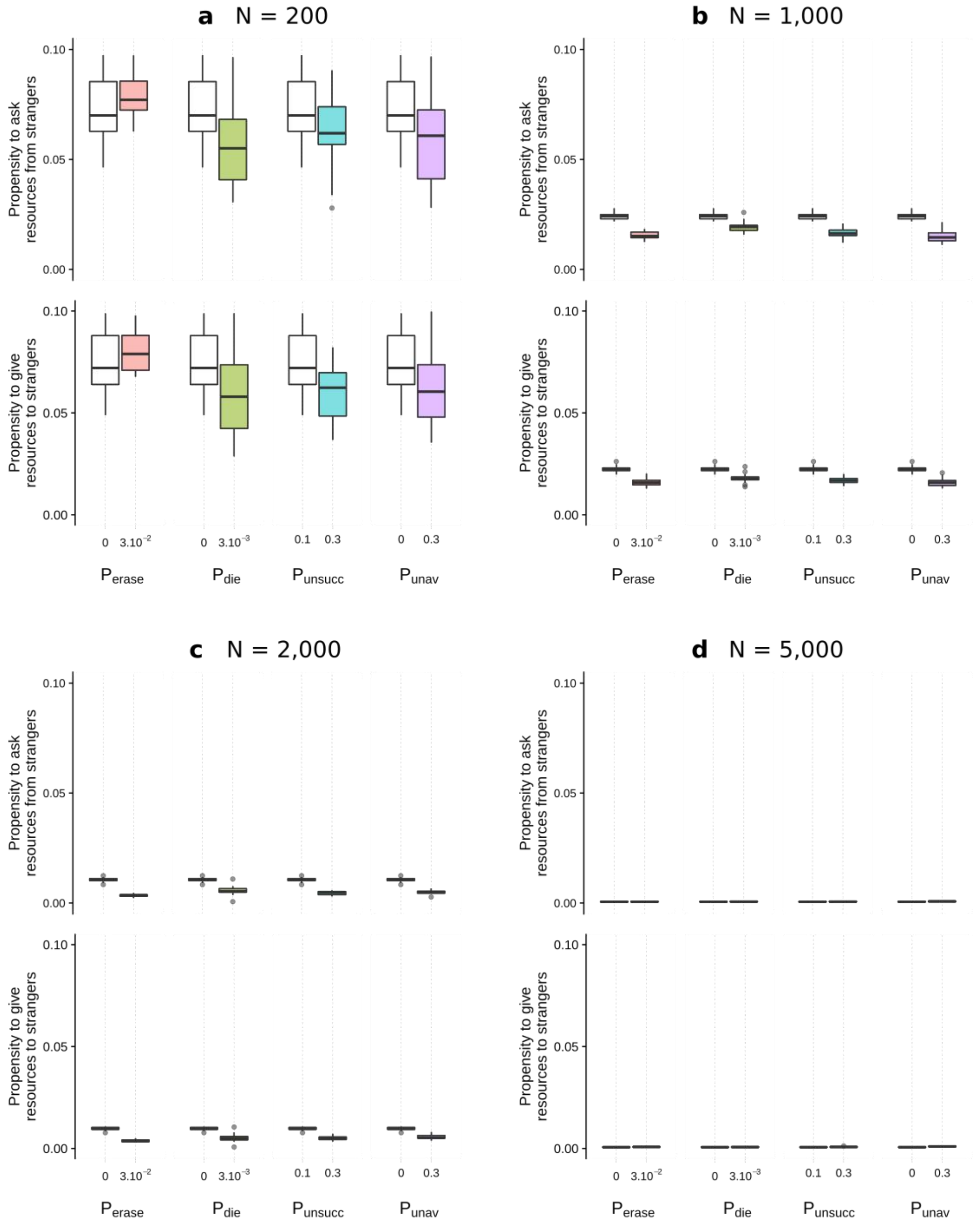

**Figure S7:** Trait values, $A_i$ and $G_i$, at evolutionary equilibrium reached after 5 million time steps, for different strengths of the prior. For high F, updating away from prior belief is more difficult, and individuals have to focus investment on few individuals to build partnerships, leading to low trait values $A_i$ and $G_i$. For high values of $P_{erase}$, $P_{die}$, $P_{unsucc}$ and $P_{unav}$, the evolutionary equilibrium is characterized by cooperative strategies focusing on the quality of each partnership rather than on the quantity of partnerships (low $A_i$ and low $G_i$) as in the main analysis.

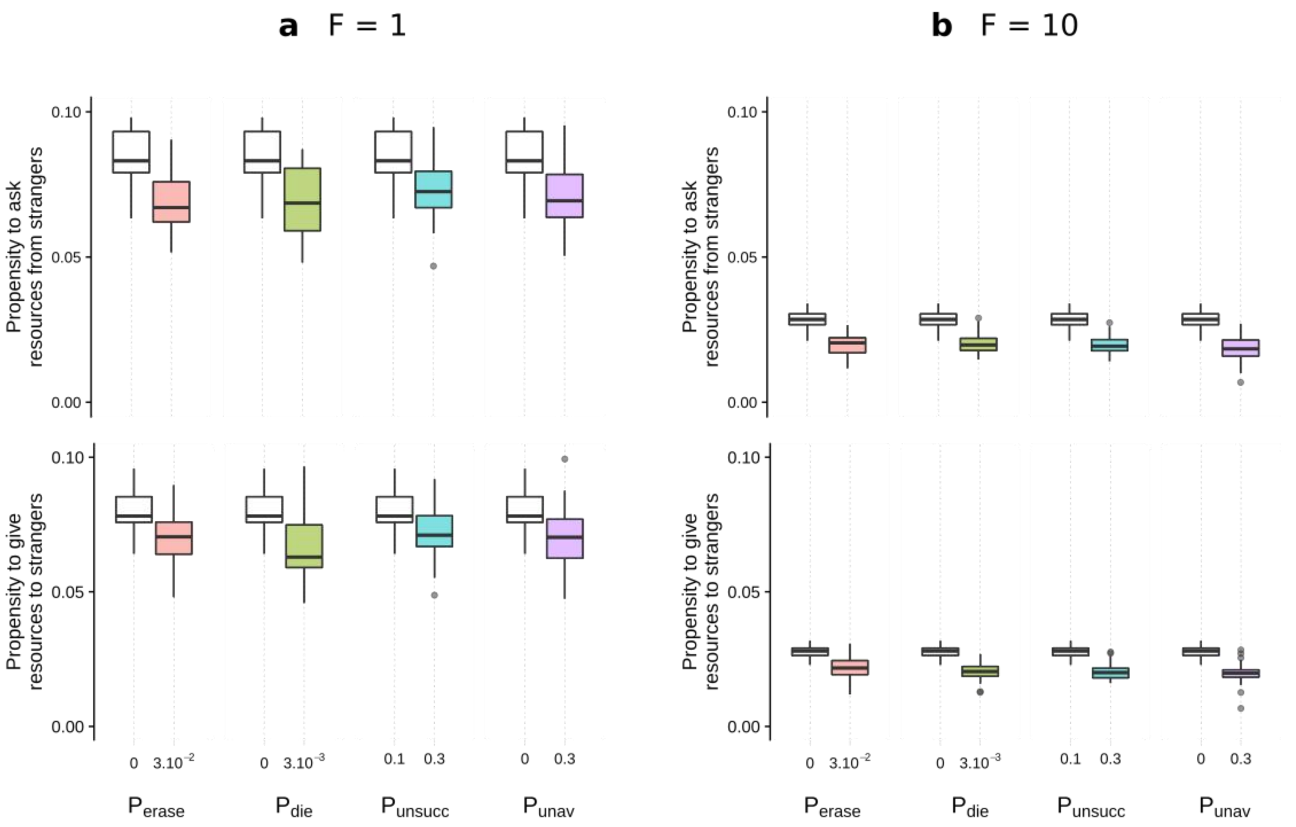

**Figure S8:** Trait values, $A_i$ and $G_i$, at evolutionary equilibrium reached after 5 million time steps, for different number of possible interactions per time step. For low values of $n_{ask}$ and $n_{give}$, individuals have less interaction opportunities, and individuals have to focus cooperative investments on few individuals to build partnerships, leading to low trait values $A_i$ and $G_i$. For high values of $P_{erase}$, $P_{die}$, $P_{unsucc}$ and $P_{unav}$, the evolutionary equilibrium is characterized by cooperative strategies focusing on the quality of each partnership rather than on the quantity of partnerships (low $A_i$ and low $G_i$) as in the main analysis.

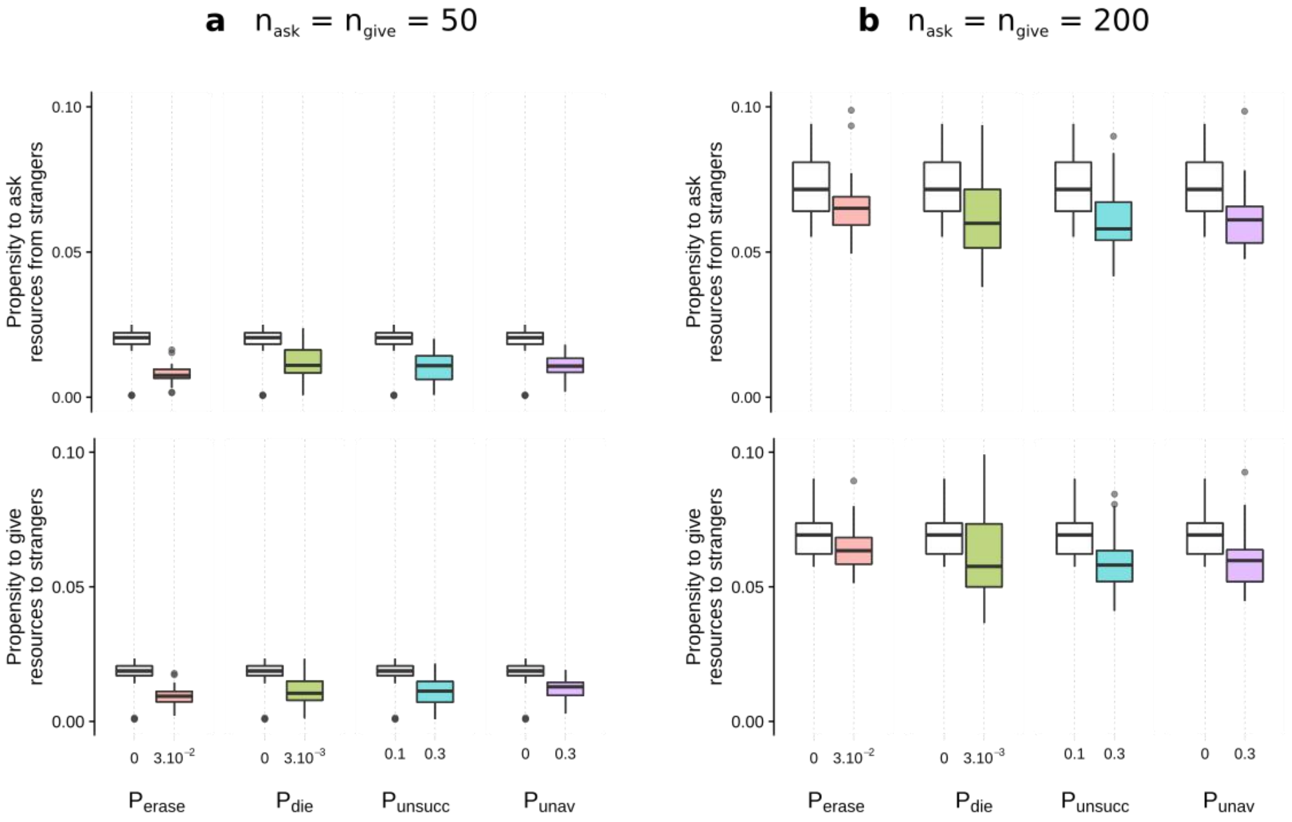

**Figure S9:** Trait values, $A_i$ and $G_i$, at evolutionary equilibrium reached after 5 million time steps, for different resource-dependent mortality functions shown in Fig. 2. For high resource-dependent mortality, individuals have to focus investment on few individuals to build partnerships, leading to low trait values $A_i$ and $G_i$. For high values of $P_{erase}$, $P_{die}$, $P_{unsucc}$ and $P_{unav}$, the evolutionary equilibrium is characterized by cooperative strategies focusing on the quality of each partnership rather than on the quantity of partnerships (low $A_i$ and low $G_i$) as in the main analysis.

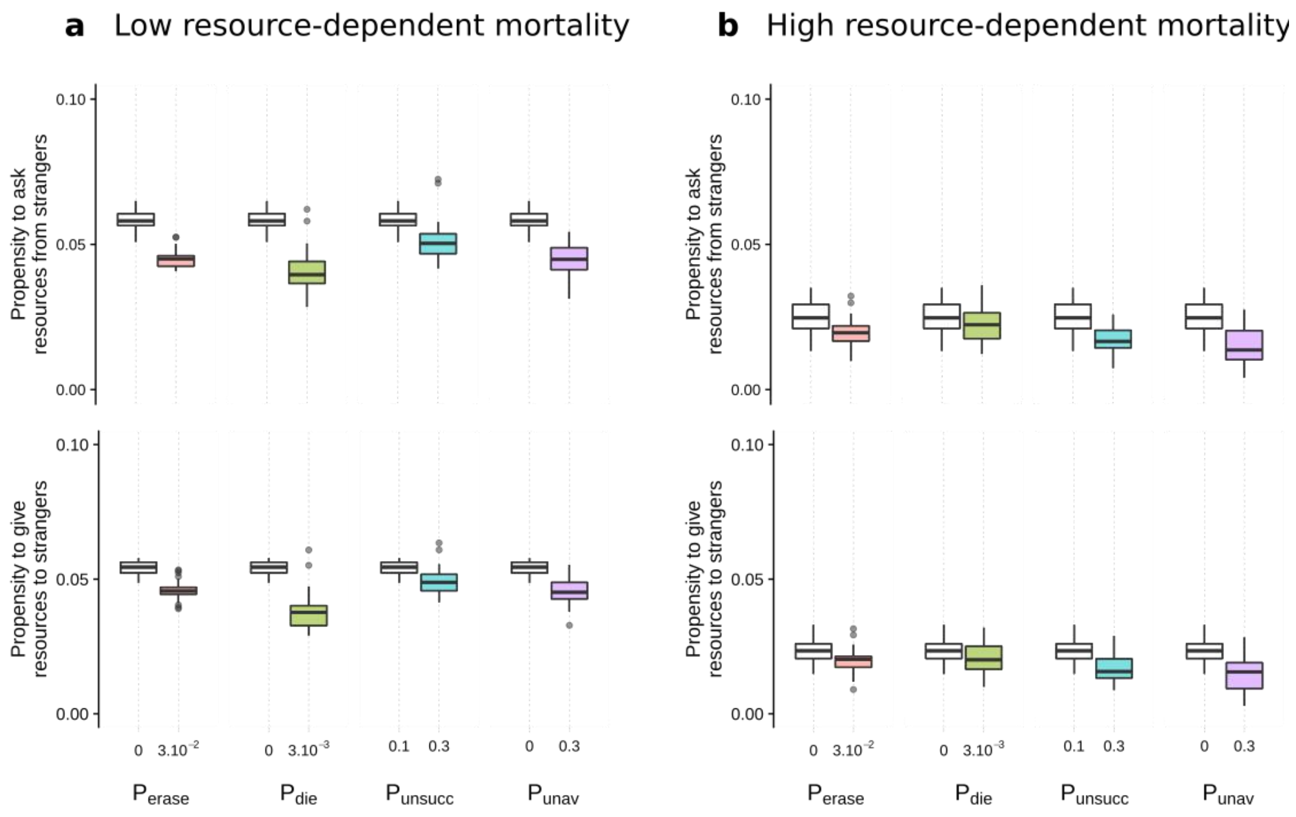

**Figure S10:** Probability of survival of unsuccessful individuals depending on the amount of resources received with a different resource-dependent mortality function (a), and trait values, $A_i$ and $G_i$, at evolutionary equilibrium reached after 5 million time steps (b). Here, we implement the survivorship function as: $P_{starv}(r) = (1 - \gamma_s)(1 - exp(-(1-r)/R_s))$, with $\gamma_s$=0.9 and $R_s$=0.4.
For high values of $P_{erase}$, $P_{die}$, $P_{unsucc}$ and $P_{unav}$, the evolutionary equilibrium is characterized by cooperative strategies focusing on the quality of each partnership rather than on the quantity of partnerships (low $A_i$ and low $G_i$) as in the main analysis.

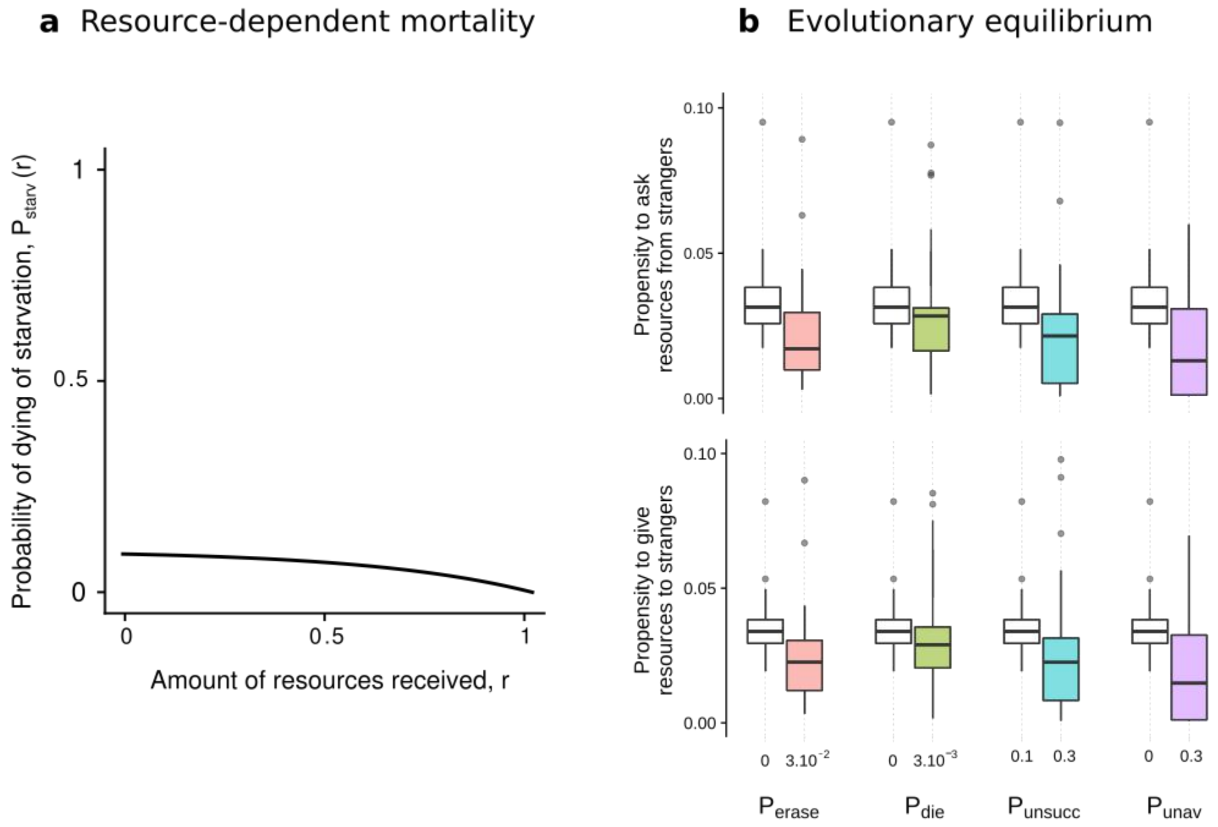

**Figure S11:** Comparison between different age measures. In the analysis, we measure age based on time steps at unsuccessful state. Here, we show that this age measure reflects age based on all time steps, such that ‘Age based on time steps at unsuccessful state’ ≈ $P_{unsucc}$ x ‘Age based on all time steps’.

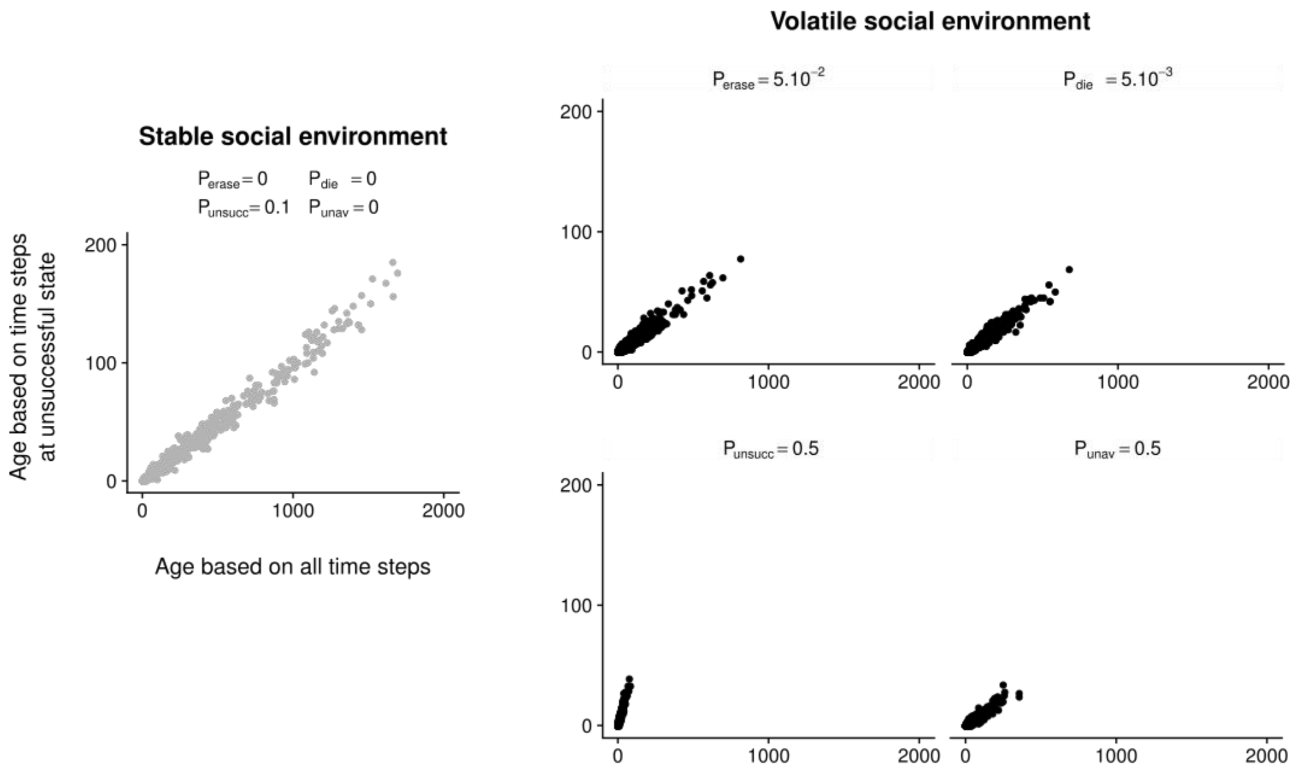

**Figure S12:** Characteristics of social relationships and survivorship of one individual i depending on its age and on its trait value in a population where $A_j = G_j = 0.035$ for all other individuals j either under a stable social environment or under a volatile social environment ($P_{die} = 5.10^{-3}$), assuming that newborn individuals have a stong social bond with their mother. As in Fig. S1 and S2, we assume that newborn individuals have initially strong social bonds with their mother (equivalent to social bonds acquired following 100 resource donations out of 100 requests). See caption of Figure 5 for more details. We get qualitatively the same results as in the case where newborn individuals build their social network from scratch (Fig. 5). In a volatile society, exploiting the benefits associated with the pre-existing social bonds (here with the mother) at the expense of diversifying cooperative investments increases survival at early age. This is crucial under a volatile social environment, even if this leads to a smaller network that is deleterious later in life (as in Fig. 5).

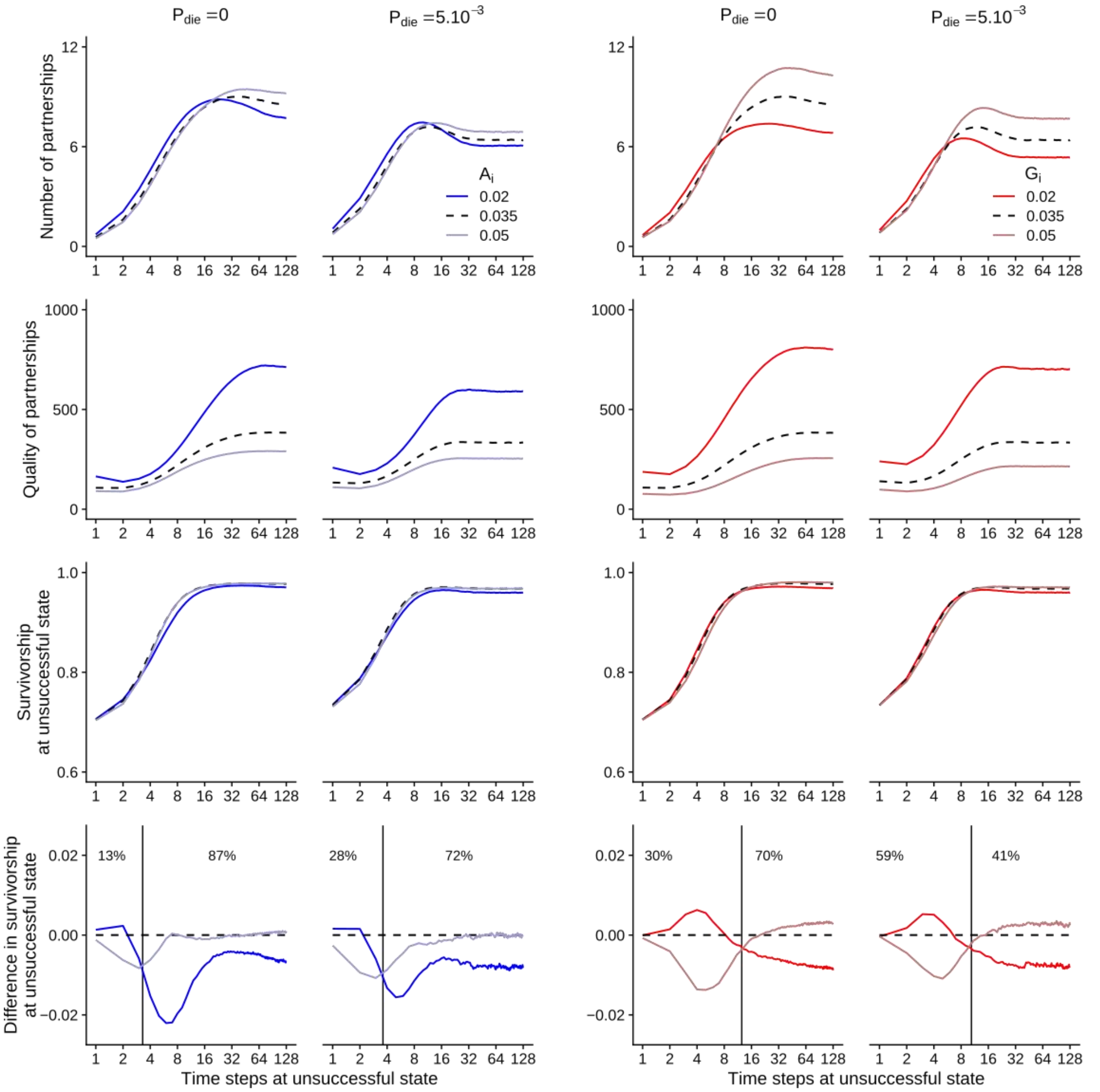

**Figure S13:** Characteristics of social relationships and survivorship of one individual i depending on its age and on its trait value in a population where $A_j = G_j = 0.035$ for all other individuals j either under a stable social environment or under a volatile social environment (**$P_{erase} = 5.10^{-2}$**). See caption of Figure 5 for more details. We get qualitatively the same results as in the case with a high probability $P_{die}$ of random death (Fig. 5) with the exception that individuals with low $A_i$ ang $G_i$ traits end up with a low number of relationships even late in life when $P_{erase} = 5.10^{-2}$ (for $A_i$=0.02 vs. for $A_i$=0.05). In a volatile society, speeding up the socializing process at an early age is crucial, even if this leads to a smaller network that is deleterious later in life (as in Fig. 5).

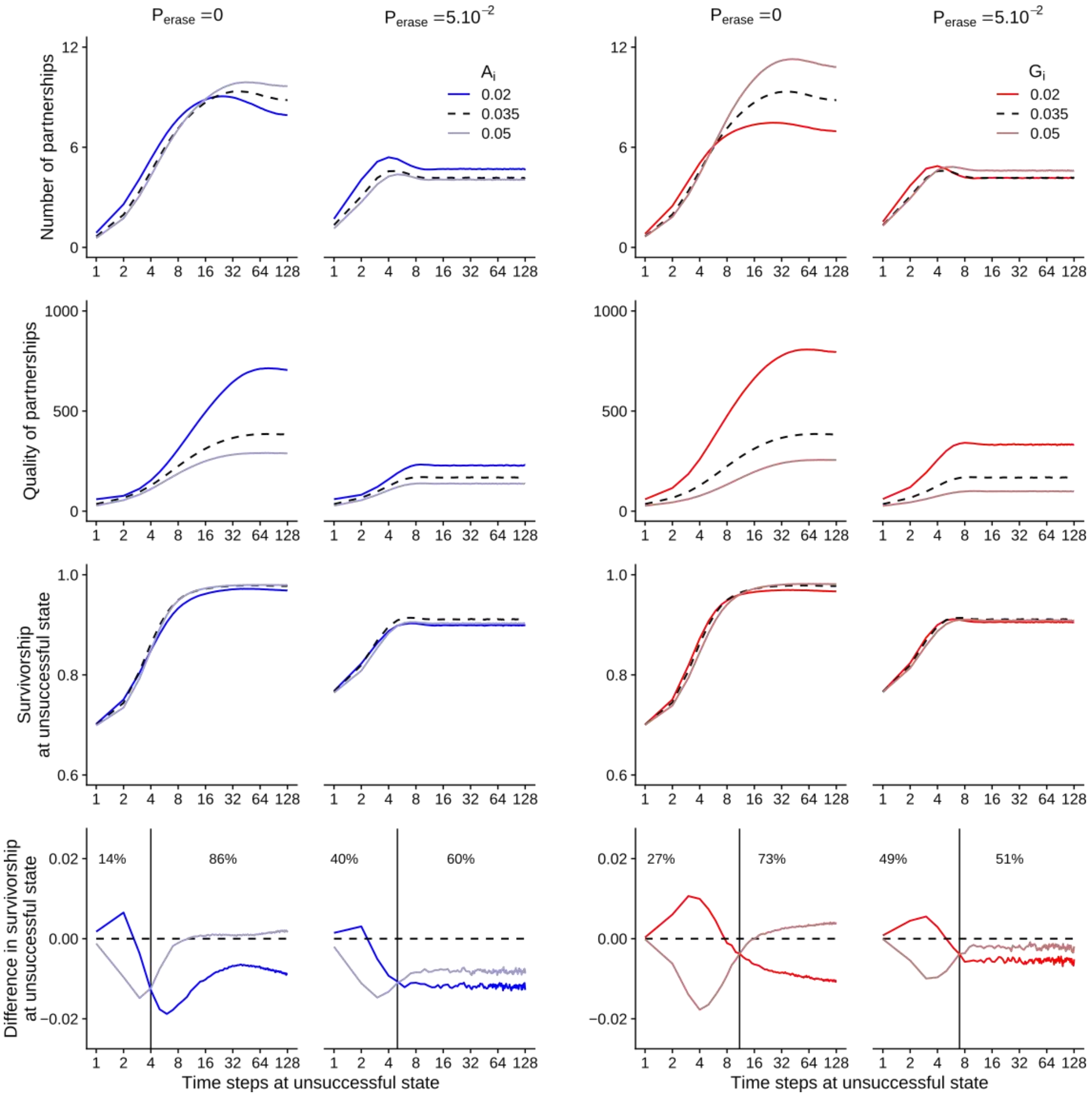

**Figure S14:** Characteristics of social relationships and survivorship of one individual i depending on its age and on its trait value in a population where $A_j$ = $G_j$ = 0.035 for all other individuals j either under a stable social environment or under a volatile social environment (**$P_{unsucc}$ = 0.5**). See caption of Figure 5 for more details. We get qualitatively the same results as in the case with a high probability $P_{die}$ of random death (Fig. 5). Early performance matters more than late performance (as in Fig. 5).

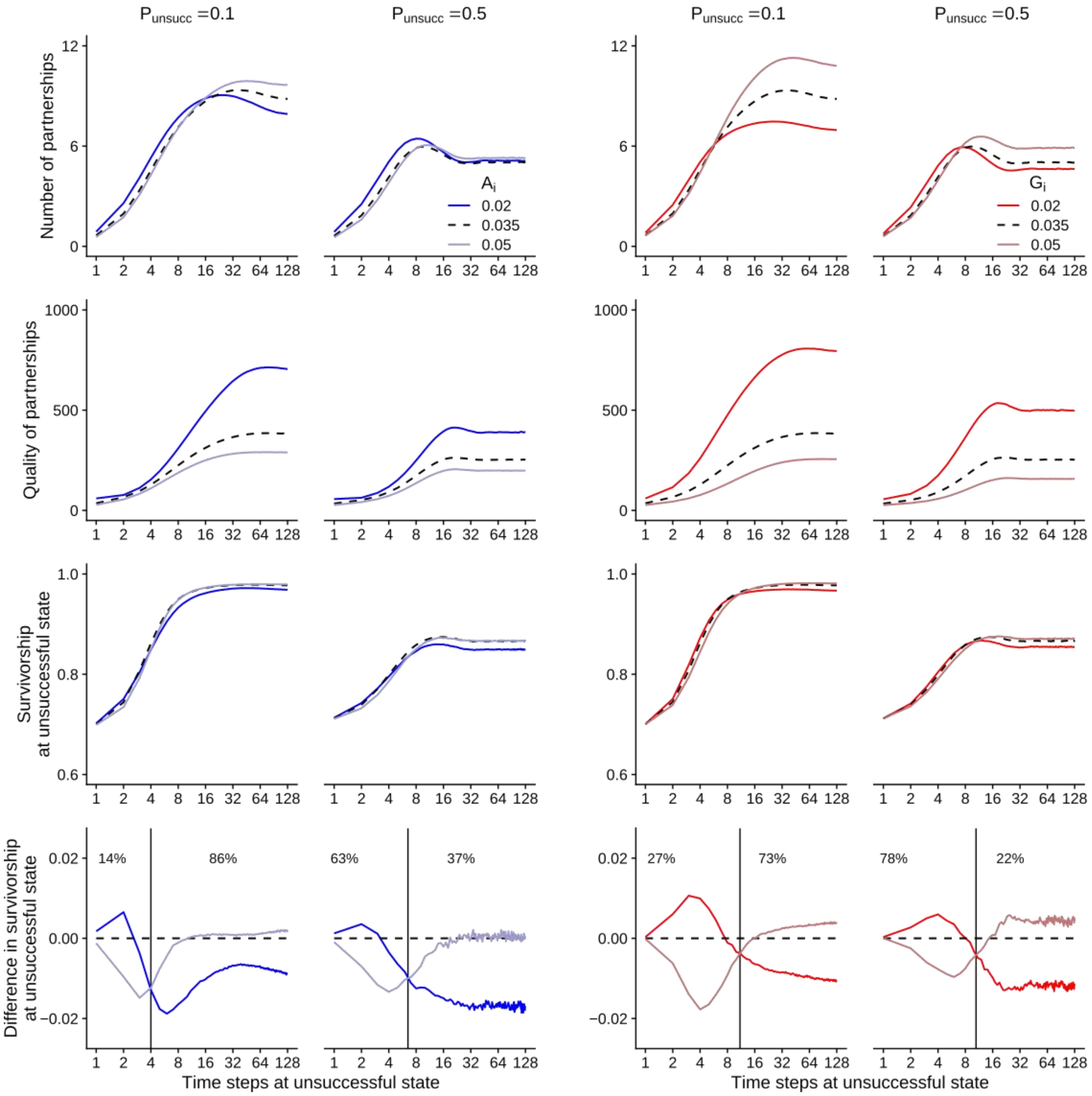

**Figure S15:** Characteristics of social relationships and survivorship of one individual i depending on its age and on its trait value in a population where $A_j$ = $G_j$ = 0.035 for all other individuals j either under a stable social environment or under a volatile social environment (**$P_{unav}$ = 0.5**). See caption of Figure 5 for more details. We get qualitatively the same results as in the case with a high probability $P_{die}$ of random death (Fig. 5). Early performance matters more than late performance (as in Fig. 5).

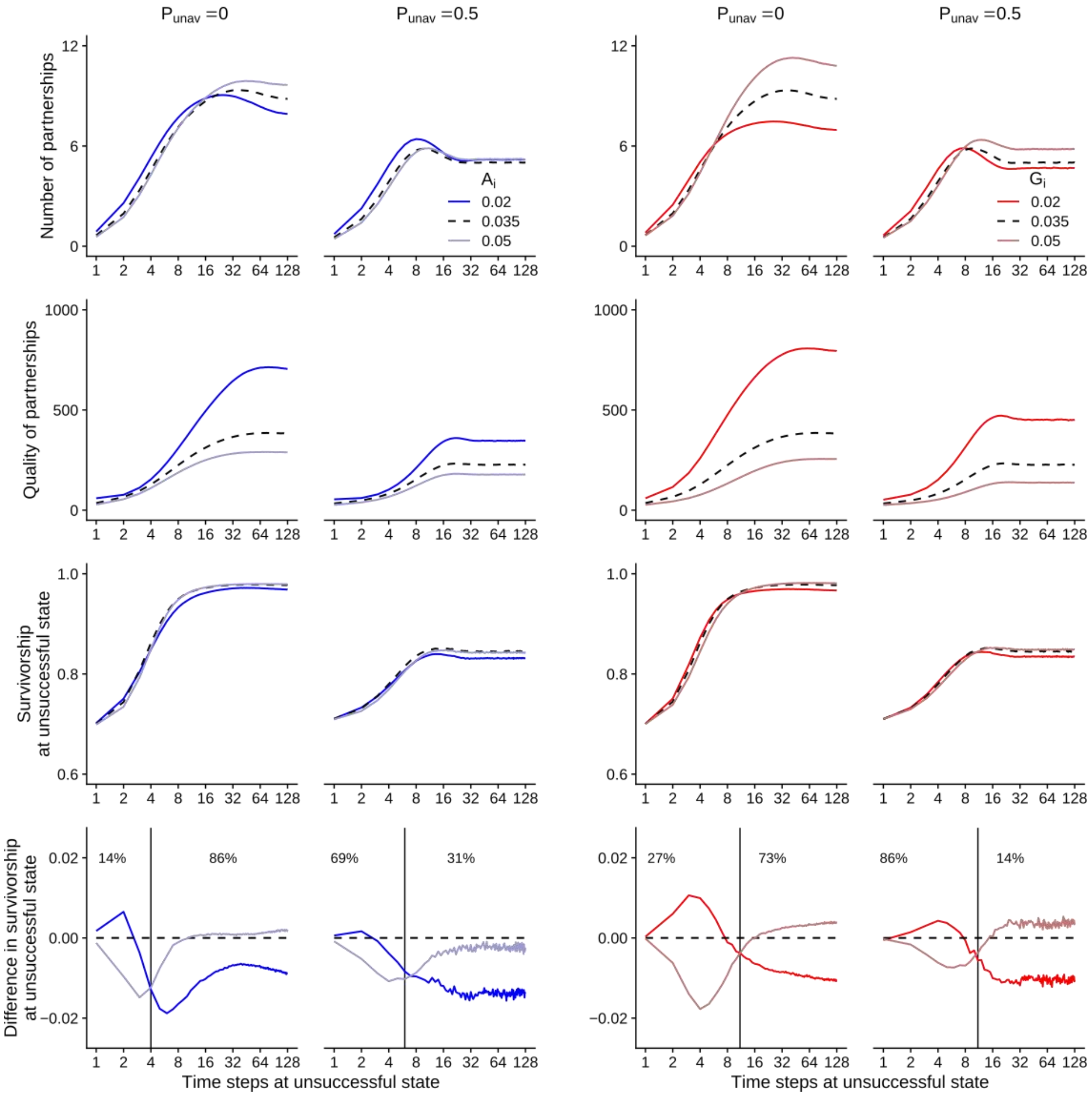

**Figure S16:** Age distribution under different social environments. A volatile social environment associates with high mortality, and therefore changes the age distribution.

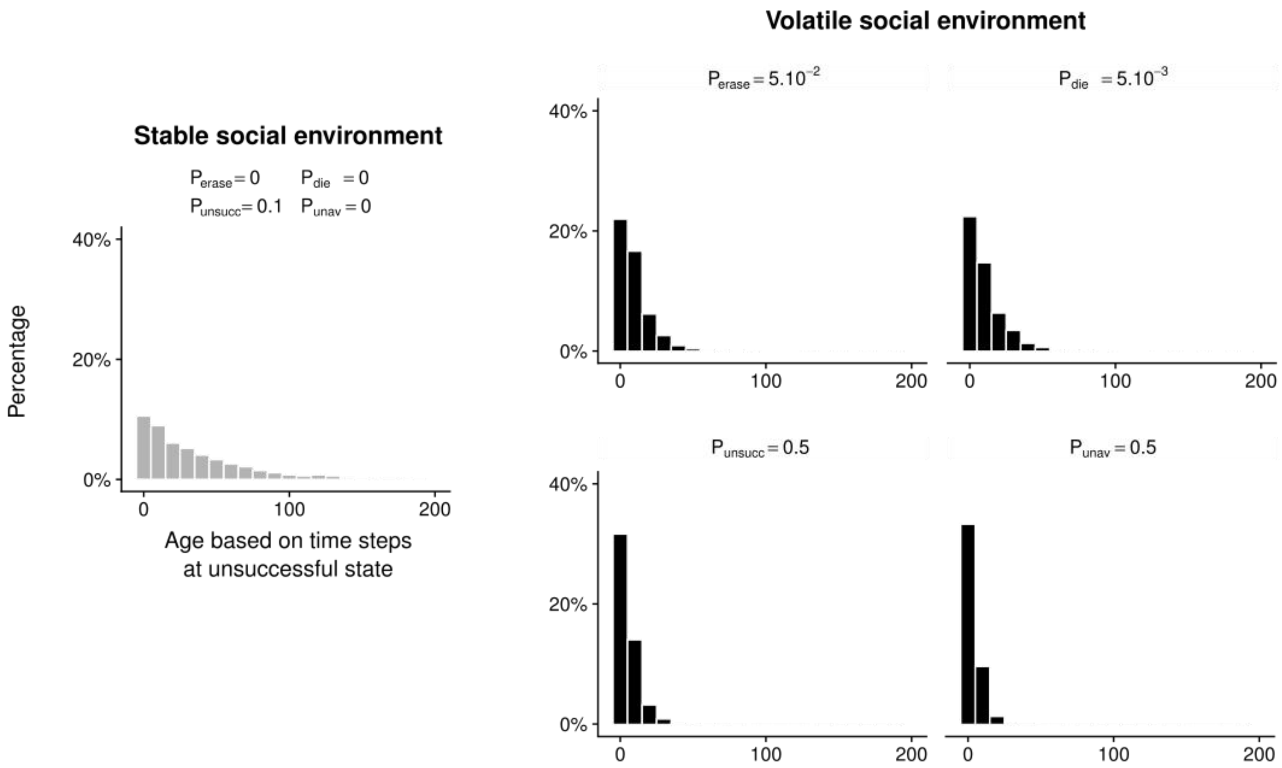
